# Sexually dimorphic effects of mutations in ccRCC driver genes in renal proximal tubule cells

**DOI:** 10.64898/2026.05.22.727114

**Authors:** Antonella Catalano, Constanze Kainz, Katharina Moos, Stefan Jäger, Marvin Müller, Yannick Federkiel, Christian Witt, Francesca Cuomo, Alex Waterhölter, Mona Schoberth, Pascal Schlosser, Anna Köttgen, Stefan Haug, Melanie Boerries, Ian J. Frew, Mojca Adlesic

**Author notes:** **Corresponding Authors and Address:** Dr. Mojca Adlesic and Prof. Dr. Ian Frew, Department of Internal Medicine I, Hematology, Oncology and Stem Cell Transplantation Medical Centre - University of Freiburg, Breisacherstraße 115, 79106 Freiburg Germany, Email addresses.

## Abstract

Clear cell renal cell carcinoma (ccRCC) arises more frequently in men than in women, but it remains unclear why this is the case. Mouse models that mimic the truncal mutation of the two most commonly affected tumor suppressor genes in human ccRCC revealed that the transcriptional effects of *Vhl* or *Vhl/Pbrm1* mutation in male and female proximal tubule cells only partially overlap. Cellular responses to mutations in these genes were also sex-specific. *Vhl* and *Vhl/Pbrm1* mutation in females, but only *Vhl/Pbrm1* mutation in males, induced the accumulation of PLIN2-coated lipid droplets. The accumulation of optically-clear lipid droplets, reflecting the hallmark clear-cell histological feature of ccRCC, only arose in *Vhl/Pbrm1* mutant male mice. Consistent with lipotoxic tubular damage, markers of tubular repair and ongoing epithelial cell proliferation correlated with the lipid accumulation phenotypes. *Vhl/Pbrm1* mutation caused stronger clonal expansion than *Vhl* mutation in males, but not in females. These findings suggest that there are likely to be sexual dimorphisms at the very earliest stages of ccRCC evolution that result from the interplay of intrinsic and mutation-induced transcriptional differences between male and female proximal tubule cells.

## INTRODUCTION

Clear cell renal cell carcinoma (ccRCC) is the most common renal malignancy and men are approximately twice as likely as women to develop this disease ^1^. The reasons for the discrepancy in tumor incidence remain unclear, however, epidemiological studies suggest that behavioral and lifestyle factors only partly contribute to the differences between the sexes ^2^. Supporting the idea that the sex imbalance in ccRCC arises in large part due to intrinsic biological factors, the renal epithelial cell-specific deletion of *Vhl*, *Trp53* and *Rb1* induced more ccRCC tumors per mouse, at earlier timepoints and with higher penetrance in male than in female mice ^3^. This suggests that at least some of the factors that predispose men to develop ccRCC more often than women are conserved in mice. ccRCC in humans and mice arises from renal proximal tubule epithelial cells ^3,4^. In this study we use a mouse genetic approach to investigate whether the enhanced propensity of men to develop ccRCC might partly arise due to intrinsic physiological differences between male and female proximal tubule cells, combined with sex-specific responses of these cells to mutations in ccRCC tumor suppressor genes.

Proximal tubule cells are the most metabolically active cell type in the kidney and are responsible for much of the kidney’s resorption of metabolites, glucose, salts and water ^5^. This transport is energetically fueled by a high mitochondrial density, which allows the generation of large amounts of ATP ^5^. Isotope-tracing studies suggest that proximal tubule cells exhibit a high degree of metabolic flexibility in terms of substrate usage to fuel the tricarboxylic acid (TCA) cycle (reviewed in ^5^). The largest contribution in the non-fasted state comes from lactate, followed by free fatty acids, glutamine, acetate and citrate, while ketone bodies can also be used under fasted conditions. Glucose normally represents a very minor source of carbon for the TCA cycle. Proximal tubule epithelial cells in mice and humans exhibit sexually dimorphic patterns of gene expression ^6–10^. Male and female proximal tubule cells express different patterns of transporters and the proximal tubules in males are responsible for a greater percentage of energy-dependent salt transport than in female kidneys, where more salt resorption occurs in the more distal tubular segments ^11,12^. Consistent with the predicted higher energetic demand necessary to fuel this transport, evidence from single cell and single nucleus RNA sequencing studies suggests that male proximal tubule cells might have higher rates of mitochondrial metabolism than female cells. Mouse proximal tubules show male-specific upregulation of genes associated with fatty acid metabolism, lipid and organic acid metabolism and oxidative phosphorylation ^6–8^, and a human study demonstrated gene set enrichment for oxidative phosphorylation, TCA cycle and electron transport chain in male versus female proximal tubules ^9^. Cultured primary human proximal tubule cells demonstrated higher levels of mitochondrial respiration in male cells than in female cells ^9^. Male mouse and human proximal tubules are more sensitive than females to ischemic injury, potentially consistent with their higher metabolic rate and mitochondrial energy dependence ^13^. There are therefore fundamental sex-specific metabolic differences in the ccRCC cell of origin. These differences are noteworthy because ccRCC is characterized by a genetically-driven, fundamental rewiring of cellular metabolism ^14^.

Approximately 90% of sporadic ccRCC tumors harbor biallelic inactivation of the von Hippel-Lindau (*VHL*) tumor suppressor gene due to loss of one copy of chromosome 3p and inactivation of the second allele by mutation, deletion or hypermethylation ^15,16^. Biallelic *VHL* inactivation is the earliest event in the process of tumor formation in the majority of sporadic and familial ccRCC cases ^17^. pVHL, the protein product of the *VHL* gene, functions as a multi-functional tumor suppressor protein that regulates numerous biochemical processes and cellular activities ^18–20^. These dysregulated processes in *VHL* mutant cells likely contribute to different aspects of tumor formation and progression. Amongst these functions, pVHL normally negatively regulates the activity of the HIF-1α and HIF-2α ^19^, ZHX2 ^21^ and SFMBT1 ^22^ transcription factors. One of the best characterized effects of loss of pVHL function is the HIF-α-mediated rewiring of cellular metabolism. *VHL* mutation in human ccRCCs induces high rates of glycolysis, reduction of mitochondrial abundance, reduction of mitochondrial electron transport, induction of reductive carboxylation of glutamine and cytoplasmic accumulation of lipids, giving rise to the eponymous clear cell appearance of ccRCC ^14^. This metabolic pattern is vastly different from the predominant mitochondrial substrate oxidation of normal proximal tubule epithelial cells. These effects of *Vhl* mutation are conserved in mouse embryonic fibroblasts, mouse primary renal epithelial cells, mouse kidneys and mouse ccRCC tumors, where unrestrained HIF-1α acts to increase glycolysis and reduce mitochondrial abundance and respiration ^23,24^.

In addition to *VHL* mutation, the evolution of familial and sporadic forms of ccRCC requires mutation in one or more tumor suppressor genes that control diverse epigenetic functions, including *PBRM1*, *BAP1*, *SETD2*, *KDM5C*, or in genes that regulate the cell cycle or cellular signaling networks, including *TP53*, *RB1*, *MTOR*, *PTEN* and *MYC* ^15,16^. *PBRM1* is the most frequently altered second-hit gene, with bi-allelic loss of function mutations arising in approximately half of all sporadic ccRCCs ^15,16^. Multi-regional DNA sequencing studies of human ccRCCs revealed that the same *PBRM1* mutation is typically found across different regions of the tumor, indicating that *PBRM1* is often mutated as an early truncal driver mutation during tumor evolution ^25^. Consistent with the human mutational data, mouse genetic studies showed that the combined deletion of *Vhl* and *Pbrm1* induces ccRCC tumors, albeit with a latency of 1-2 years after inducible gene deletion in proximal tubule epithelial cells in adult mice ^26–28^. PBRM1 (also known as BAF180) functions as a component of the SWI/SNF family PBAF chromatin remodelling complex, which controls the nucleosome structure of chromatin, regulating the accessibility of DNA for other epigenetic or transcriptional regulators. PBAF activity can either act to promote or inhibit gene transcription. It is therefore likely that the cooperative effects of mutations in *VHL* and *PBRM1* at least partly contribute to ccRCC formation through alterations in gene expression. The transcriptional consequences of loss of PBRM1 function in ccRCC appear to be highly dependent on the context of the cell or animal model used, as might be expected from a general chromatin regulator ^27,29–32^.

In the current study we developed new mouse models based on *Vhl* and *Pbrm1* deletion to address two major issues. Firstly, we aimed to characterize the earliest molecular changes in proximal tubule cells following deletion of *Vhl*, or of *Vhl* together with *Pbrm1* (*Vhl/Pbrm1*), to gain insight into how loss of PBRM1 function initially cooperates with loss of pVHL function to ultimately prime these cells to develop into ccRCC tumors. Secondly, we aimed to investigate whether *Vhl* or *Vhl/Pbrm1* deletions cause sex-specific differences in renal proximal tubule epithelial cells to ask whether different responses to the same genetic perturbations might potentially contribute to the different propensities of men and women to develop ccRCC.

## RESULTS

### Mouse models that allow genetic marking of *Vhl* and *Vhl/Pbrm1* mutant renal epithelial cells

*Ksp1.3-CreER^T^*^2^ transgenic mice allow tamoxifen-inducible Cre-mediated recombination specifically in renal epithelial cells throughout the nephron ^33^. We demonstrated that this transgene targets the specific cell type in the mouse proximal tubular epithelium that gives rise to *Vhl/Trp53/Rb1* mutant ccRCC tumors, which show many phenotypic and molecular similarities to human ccRCC ^3^, validating the utility of this Cre transgene for modelling human ccRCC in mice. With the aim of more accurately reflecting the two most frequently-arising truncal driver gene mutations in human ccRCC we first sought to reproduce published mouse studies demonstrating that combined *Vhl/Pbrm1* mutation in adult kidneys can induce ccRCC evolution after a long latency ^26–28^. We fed i) *Ksp1.3-CreER^T^*^2^, ii) *Ksp1.3-CreER^T^*^2^*; Vhl^fl/fl^* and iii) *Ksp1.3-CreER^T^*^2^*; Vhl^fl/fl^; Pbrm1^fl/fl^*mice with tamoxifen-containing food for two weeks and followed the mice using regular ultrasound imaging to detect the formation of renal cysts, which can represent a precursor of some ccRCCs in humans and mice ^3,26,34–36^ (Supplementary Fig. 1a). *Vhl* deletion was sufficient to induce cyst formation in about half of all male mice and this was weakly enhanced by additional *Pbrm1* mutation. In female mice, *Vhl/Pbrm1* mutation induced cysts in half of the mice, while only 10% of *Vhl* mutant mice showed cysts. Histological analyses of 20 mice of each genotype and sex harvested 25 months after tamoxifen feeding confirmed the presence of cysts at frequencies that reflected the ultrasound imaging. While no tumors developed in *Vhl* mutant mice, 16% of male and 22% of female mice developed ccRCCs (Supplementary Fig. 1b,c). We conclude that while combined *Vhl* and *Pbrm1* mutation using this experimental model is not a strong driver of tumor formation, it nonetheless does lead to ccRCC development and appears to reflect the decades-long process of human ccRCC evolution ^37^.

To study the earliest consequences of *Vhl* and *Pbrm1* mutation prior to tumor initiation we crossed the *Ubow* transgene ^38^ into the above-described mouse strains. *Ubow* mice act as a Cre-activity reporter in which the ubiquitously expressed *Ubc* promoter drives expression of the *Brainbow1* construct, a multi-colored fluorescent transgene encoding constitutive expression of dTomato (red). Upon Cre-mediated recombination, cells lose dTomato and gain expression of either CFP (cyan) or YFP (yellow). This switch is definitive and is passed on to cellular progeny, allowing clonal tracking of all cells that were exposed to functional Cre activity. We generated i) *Ksp1.3-CreER^T^*^2^*; Ubow*^Tg/Tg^, ii) *Ksp1.3-CreER^T^*^2^*; Vhl^fl/fl^; Ubow*^Tg/Tg^ and iii) *Ksp1.3-CreER^T^*^2^*; Vhl^fl/fl^; Pbrm1^fl/fl^; Ubow*^Tg/Tg^ mice (Fig. 1a). Feeding for two weeks with tamoxifen-containing food generated fluorescently marked renal epithelial cells that were either wild type (hereafter referred to as WT), *Vhl* mutant (hereafter referred to as *Vhl*) or *Vhl/Pbrm1* mutant (hereafter referred to as *Vhl/Pbrm1*). Immunohistochemical staining using an anti-GFP antibody that recognises both CFP and YFP, but not dTomato, confirmed the expected stochastic pattern of induced Cre activity throughout the kidney (Fig. 1b) and within individual tubules (Fig. 1c). Littermate mice lacking Cre showed no staining, verifying the specificity of the antibody for detection of Cre-recombined clones (Supplementary Fig. 2a). *Vhl* deletion was validated by upregulation of the HIF-1α target protein CA9 specifically in clonally-marked (CFP/YFP positive) proximal tubule cells (lotus tetragonolobus lectin coupled to FITC (LTL-FITC) positive) (Fig. 1d). PBRM1 immunoreactivity was lost in clonally marked (CFP/YFP positive) proximal tubule (MEGALIN positive) cells (Fig. 1e, Supplementary Fig. 2b). These analyses validated that the mouse models induce deletions of the floxed genes in clonally-marked proximal tubule cells.

**Fig. 1.**
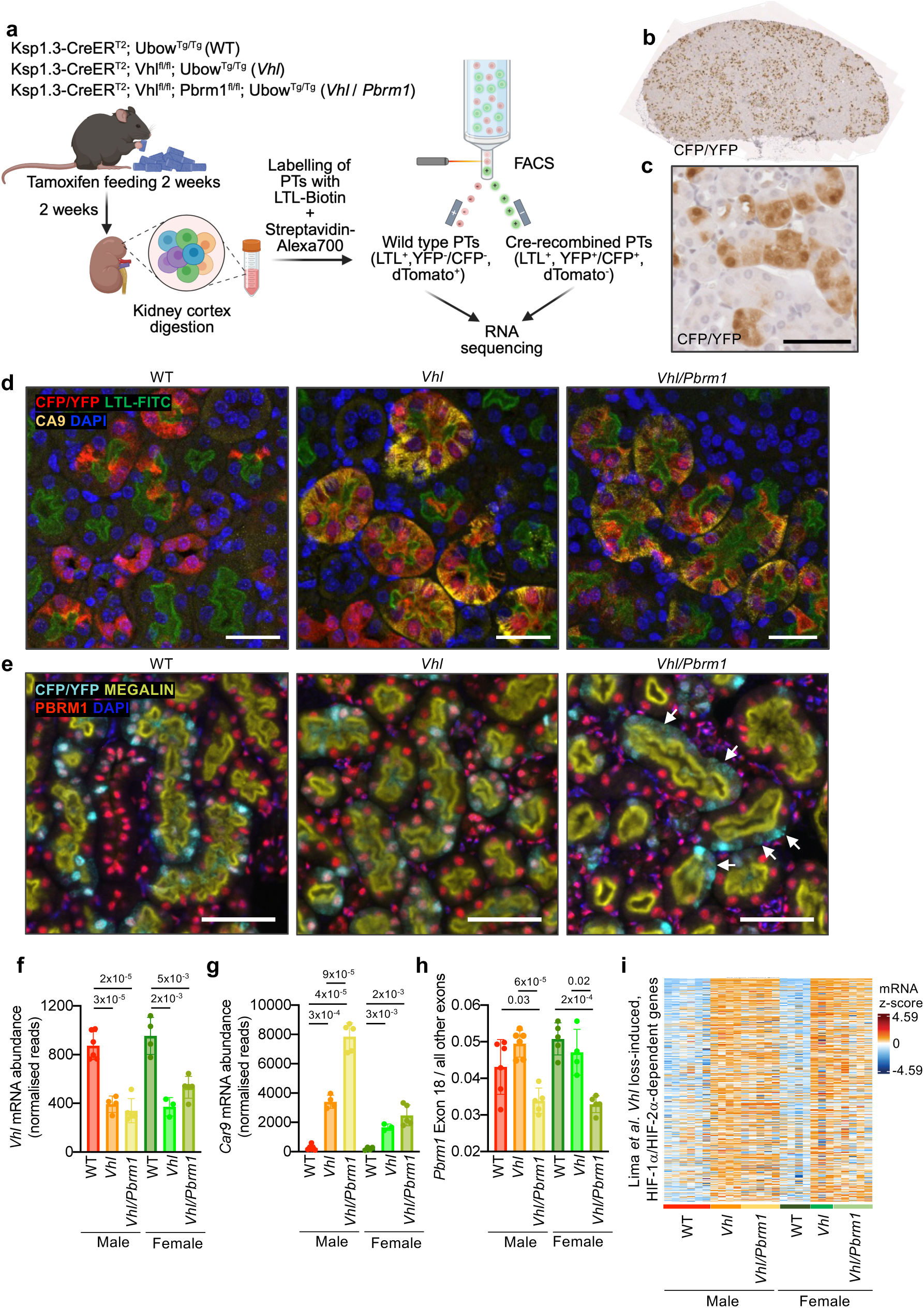
Genetic labelling of *Vhl* and *Vhl/Pbrm1* mutant renal epithelial cells. **a** Overview of the experimental system and sorting strategy used for RNA-seq of proximal tubule cells. **b,c** Representative immunohistochemical staining of tamoxifen-fed kidneys using an anti-GFP antibody that recognises CFP and YFP. Scale bar = 20 μm. **d** Immunofluorescence staining of WT, *Vhl* and *Vhl/Pbrm1* kidneys showing Cre-marked cells (anti-CFP/YFP), proximal tubules (LTL-FITC), the HIF-α target CA9 (anti-CA9) and nuclei (DAPI). Scale bar = 50 μm. **e** Immunofluorescence staining of WT, *Vhl* and *Vhl/Pbrm1* kidneys showing Cre-marked cells (anti-CFP/YFP), proximal tubules (anti-MEGALIN), PBRM1 (anti-PBRM1) and nuclei (DAPI). Arrows highlight clones in the *Vhl/Pbrm1* kidney. Scale bar = 50 μm. Split channel zoomed images are shown in Supplementary Fig. 2. **f-h** Relative mRNA abundance (normalised reads from RNA-seq) of *Vhl* (**f**), *Car9* (**g**) and ratio of *Pbrm1* floxed exon 18 to all other *Pbrm1* exons (**h**). Data represents mean ± std. dev. of 4-6 mice (represented as dots) of the indicated genotype and sex. *P* values were calculated by two-sided Student’s t-test without adjustment for multiple comparisons. **i** Gene expression heatmap of 467 previously-identified HIF-α target genes ^40^. Each column represents an independent mouse of the indicated genotype and sex. Gene lists and their expression values are provided in the Source Data Table.

### Sex-specific effects of *Vhl* and *Vhl/Pbrm1* deletion on gene expression

To investigate how the earliest genetic mutations that arise in ccRCC affect proximal tubules in male and female mice we used flow cytometry to isolate proximal tubule cells from male and female kidneys of each genotype and sex (n = 4-6 mice) two weeks after the end of a two-week period of tamoxifen feeding. Kidney cortices were digested to single cell suspensions and sorted based on positivity for the proximal tubule marker LTL and either i) positivity for dTomato and negativity for CFP/YFP (absence of Cre activity) or ii) negativity for dTomato and positivity for CFP/YFP (presence of Cre activity) (Fig. 1a, Supplementary Fig. 3a). RNA-seq of the sorted cells allowed determination of the transcriptomes of WT, *Vhl* and *Vhl/Pbrm1* mutant male and female renal proximal tubule epithelial cells between 2 and 4 weeks after gene deletion. Normalized RNA-seq data of dataset i) are provided in Supplementary Data Table 1 and of dataset ii) in Supplementary Data Table 2. Analyses of the RNA-seq datasets using the CellMatchR algorithm that correlates kidney cell type-specific RNA expression patterns to bulk RNA-seq samples ^39^ revealed that all samples had a high correlation to specific gene expression signatures of S1 and S2 proximal tubule cells, and a slightly lower correlation to S3, validating the efficiency of cell sorting (Supplementary Fig. 3b-d). Principal component analyses revealed separation of the samples based on sex (Supplementary Fig. 4a) and on genotype (Supplementary Fig. 4b). Cre-mediated deletion of *Vhl* in the *Vhl* and *Vhl/Pbrm1* mutant genotypes was shown by decreased mRNA abundance of *Vhl* (Fig. 1f) and increased mRNA abundance of the HIF-1α target gene *Car9* (Fig. 1g). Decreased abundance of mRNA reads corresponding to the floxed exon 18 of *Pbrm1* compared to reads from all other non-floxed exons of *Pbrm1* confirmed the Cre-mediated mutation of *Pbrm1* specifically in the *Vhl/Pbrm1* mutant genotype (Fig. 1h). We next probed the expression levels of HIF-1α- or HIF-2α-dependent genes that were previously shown to be upregulated in single cell RNA sequencing experiments of mouse proximal tubule epithelial cells following *Vhl* deletion ^40^. Upregulation of these genes in male and female *Vhl* and *Vhl/Pbrm1* mutant cells provided an independent validation of functional *Vhl* gene deletion in our model and demonstrated that *Pbrm1* co-mutation does not have a large impact on the majority of HIF-α target genes that are induced by *Vhl* loss (Fig. 1i).

We first compared the transcriptomes of male and female wild type proximal tubule cells that had not experienced Cre activity. To externally validate our dataset, we examined the expression levels of genes that were previously found to be sex-specifically expressed in S1, S2 or S3 proximal tubules in a single nucleus RNA sequencing experiment ^6^, revealing a good overlap in the detection of male and female enriched genes (Supplementary Fig. 4c). In our data we identified 168 genes whose expression was more than 2-fold higher (*p*_adj_ < 0.05) in male cells and 119 genes whose expression was more than 2-fold higher in female cells (Fig. 2a). Probing the expression of the top male- and female-specific genes from our dataset (Supplementary Fig. 4d) using scRNA-seq data from the Kidney Cell Explorer ^8^ showed that these genes are expressed in the expected sex-specific manner in one or more of the S1, S2 or S3 proximal tubule segments, providing an additional level of validation of our dataset.

**Fig. 2.**
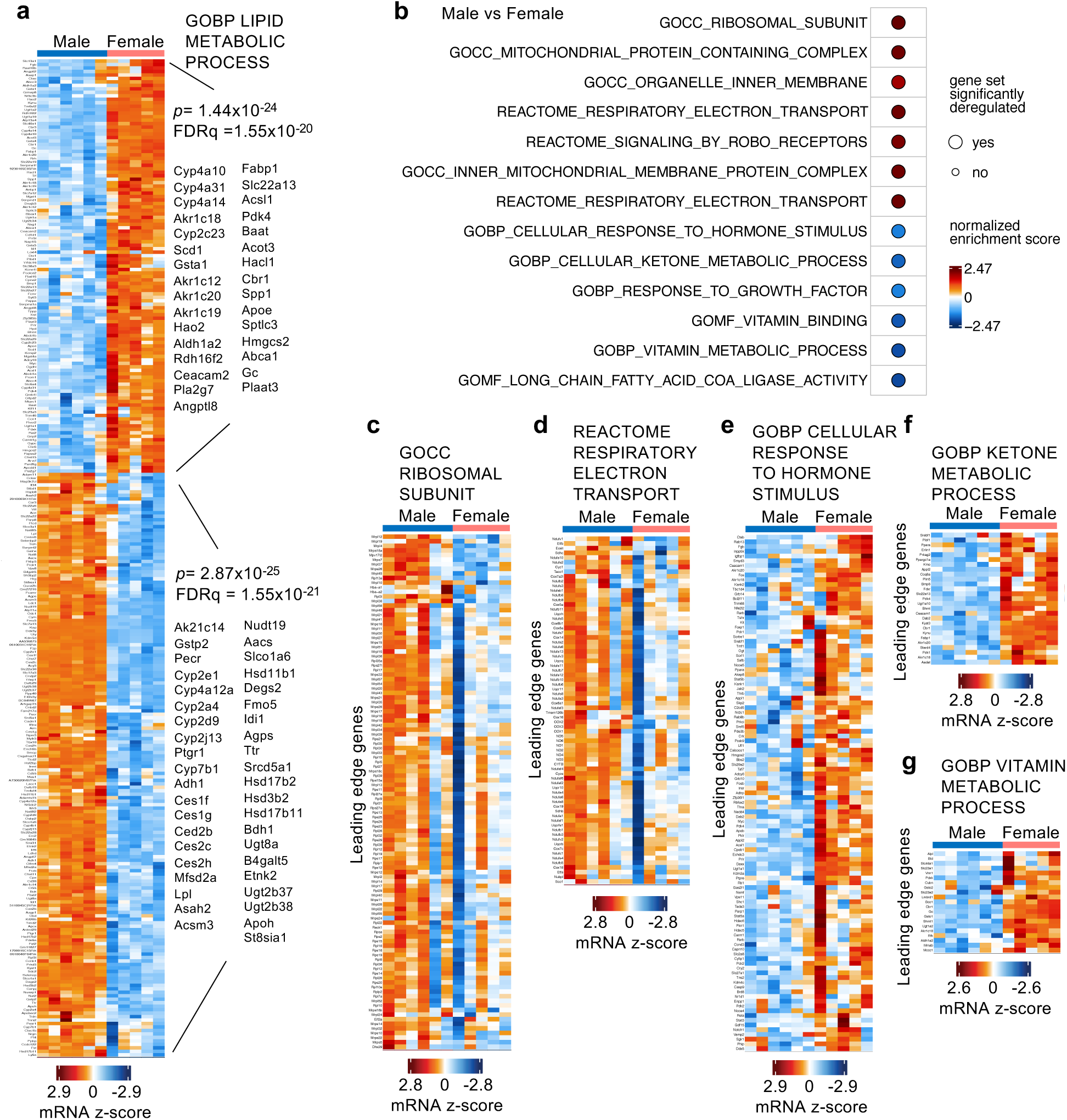
Sex-specific gene expression in proximal tubule cells. **a** Gene expression heatmap of sex-specific differentially expressed genes in wild type proximal tubule cells. GSEA scores for the term GOBP Lipid Metabolic Processes and lists of genes in each sex that are enriched in this GSEA term are shown. **b** Selected differentially enriched GSEA terms between male and female proximal tubule epithelial cells. **c-g** Gene expression heatmaps of the leading edge genes that drive the enrichment of the indicated GSEA scores. Each column represents proximal tubules derived from independent male or female mice. For all heatmaps the gene lists and their expression values are provided in the Source Data Table.

The set of sex-specific, differentially expressed genes contained many genes that belong to the GOBP term „Lipid metabolic process“in both sexes (Fig. 2a), but different genes that belong to this term were upregulated in either male or female cells. These differentially expressed genes are suggestive of different metabolic modes in males and females. Male cells show higher expression of genes that are involved in lipoprotein or fatty acid uptake (*Slco1a6, Apoh, Ttr, Mfsd2a*), oxidative metabolism of fatty acids (*Pecr, Cyp4a12a, Acsm3*), ketone utilization (*Bdh1*), lipid oxidation (*Cyp2e1, Cyp4a12a, Cyp2a4, Cyp2d9, Cyp2j13, Cyp7b1*), steroid interconversion and cholesterol biosynthesis (*Hsd11b1, Hsd17b2, Hsd17b11, Hsd3b2, Srd5a1, Cyp7b1, Idi1, Akr1c14*), sphingolipid metabolism (*Asah2, Degs2, St8sia1, B4galt5, Etnk2, Agps*) and enzymes involved in detoxification of lipids and xenobiotics (*Ugt2b37, Ugt2b38, Ugt8a, Ces1f, Ces1g, Ces2b, Ces2c, Ces2h, Gstp2, Ptgr1, Fmo5*). In contrast, female cells show higher expression of genes involved in fatty acid synthesis (*Scd1, Acsl1, Acot3, Hacl1, Sptlc3*), higher expression of *Hmgcs2*, encoding the rate limiting protein for ketone body synthesis, genes involved in promoting mitochondrial and peroxisomal fatty acid oxidation (*Pdk4, Hao2*), lipid and lipoprotein transport (*Apoe, Abca1, Fabp1, Gc*), aldehyde metabolism and detoxification of lipid peroxidation (*Aldh1a2, Akr1c12, Akr1c19, Akr1c20, Rdh16, Cbr1, Gsta1*). In conclusion, the transcriptome of male cells suggests a strong fatty-acid-oxidizing, steroid-processing, detoxification-active metabolic phenotype, while female cells are predicted to balance fatty acid oxidation and synthesis and engage in ketogenesis.

Gene set enrichment analyses (GSEA) of the entire transcriptome (Fig. 2b, Supplementary Data Table 3) further revealed that male proximal tubule cells show upregulation of terms associated with ribosomal biogenesis (Fig. 2c), suggestive of a high rate of protein synthesis. Numerous nuclear-encoded respiratory electron transport genes encompassing all mitochondrial complexes I-V as well as regulators of, or assembly factors for, electron transport chain complexes were also highly expressed in male cells (Fig. 2d), consistent with the idea of high mitochondrial content and activity of male proximal tubule epithelial cells. Female cells show enrichment of a broad GSEA term associated with cellular response to hormones (Fig. 2e), encompassing genes that regulate metabolism, energy homeostasis, insulin signaling and transcriptional and epigenetic regulators. GSEA terms related to ketone metabolic processes (Fig. 2f) and vitamin metabolic processes (Fig. 2g) are also enriched in female cells.

We conclude that the normal transcriptome of the ccRCC cell of origin differs between males and females and that pathways involved in different aspects of metabolism, mitochondrial energy generation and protein translation are likely to be differently active in the two sexes.

We next asked whether *Vhl* or *Vhl/Pbrm1* mutation might also differently affect gene transcription in male and female cells. Deletion of *Vhl* or *Vhl/Pbrm1* did not broadly alter the expression of the above-described intrinsically sex-specific genes (Supplementary Fig. 4e). However, differential gene expression analyses revealed strong sex-specific transcriptomic effects of the introduced mutations. Approximately half of all genes that were upregulated (logfold change (logfc) > 1, *p*_adj_< 0.05) in *Vhl* null cells were shared and half were male- or female-specific (Fig. 3a). Only very few genes were commonly downregulated (logfc <-1, *p*_adj_< 0.05) in male and female *Vhl* null cells (Fig. 3a). *Vhl/Pbrm1* deletion caused both the up- and downregulation of more genes in male cells than in female cells, with 65% of all upregulated genes in female cells being shared with male cells and approximately 50% of all downregulated genes in female cells being shared with male cells (Fig. 3b). More genes were up- or downregulated in *Vhl/Pbrm1* mutant cells compared to *Vhl* mutant cells from males than from females, with 55% of the female-upregulated, *Pbrm1*-dependent genes and 42% of the female-downregulated, *Pbrm1*-dependent genes, being shared with males (Fig. 3c). Heatmap visualisation of differentially expressed genes revealed the complex interplay between gene mutations and sex (Fig. 3d). Notable are the sets of genes that are only downregulated in *Vhl/Pbrm1* mutant male but not female cells, as well as genes that are upregulated by *Vhl* and/or *Vhl/Pbrm1* mutation only in male cells. These analyses demonstrate that *Pbrm1* mutation modulates the transcriptional response to *Vhl* deletion, that it does so more strongly in male than female cells and that the transcriptional consequences of *Vhl* or *Vhl/Pbrm1* mutation in proximal tubule cells only partially overlap between males and females.

**Fig. 3.**
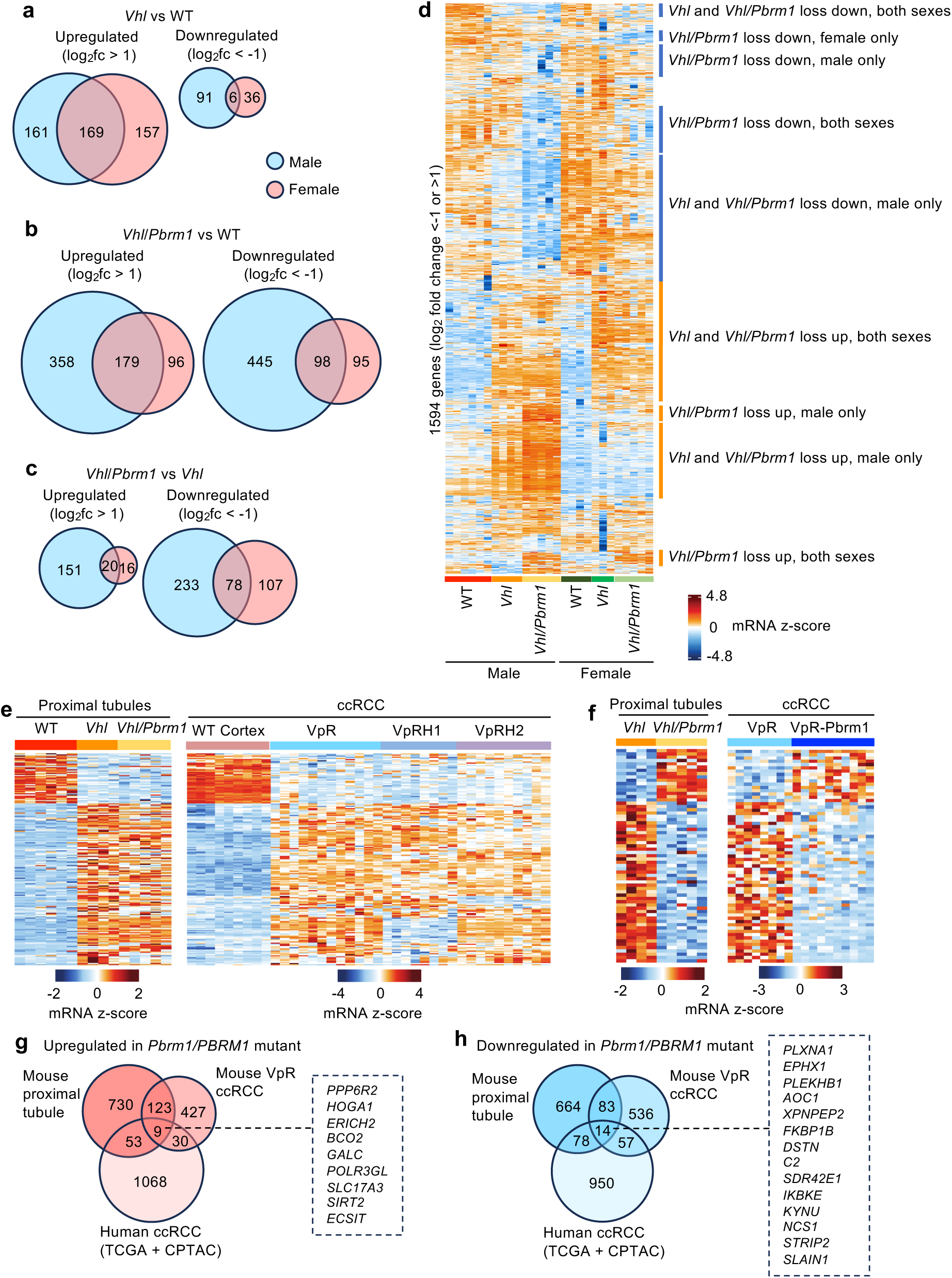
*Vhl* and *Vhl/Pbrm1* mutations induce ccRCC-relevant, sex-specific gene expression patterns in proximal tubules. **a-c** Venn diagrams showing overlapping differentially expressed genes (log2 fold change >1 or <-1, *p*adj < 0.05) between male and female mice in the comparisons of *Vhl* vs WT (**a**), *Vhl/Pbrm1* vs WT (**b**) and *Vhl/Pbrm1* vs *Vhl* (**c**). **d** Gene expression heatmap of sex-specific differentially expressed genes induced by *Vhl* and *Vhl/Pbrm1* mutations. **e** Gene expression heatmap of genes that are altered by *Vhl* or *Vhl/Pbrm1* mutation in male proximal tubule cells (left heatmap) and also differentially expressed in the comparison of normal kidney cortex to ccRCCs derived from *Vhl/Trp53/Rb1* mutant (VpR), *Vhl/Trp53/Rb1/Hif1a* mutant (VpRH1) or *Vhl/Trp53/Rb1/Hif2a* mutant (VpRH2) male mice. **f** Gene expression heatmap of genes that are only altered due to *Vhl/Pbrm1* mutation in male proximal tubule cells (left heatmap) and are also differentially expressed in the comparison of *Vhl/Trp53/Rb1* mutant (VpR) and *Vhl/Trp53/Rb1/Pbrm1* mutant (VpR-Pbrm1) male mouse ccRCCs. Each column in all heatmaps represents proximal tubules derived from independent mice of the indicated genotype and sex or of the indicated normal kidney or ccRCC tumor genotype. For all heatmaps in d-f the gene lists and their expression values are provided in the Source Data Table. **g,h** Venn diagram showing overlapping PBRM1-dependent up- (**g**) and downregulated (**h**) genes in mouse proximal tubule cells, mouse VpR ccRCC and human ccRCC. The human homologues of the genes that are commonly altered in all three settings are shown in the dotted boxes.

### Transcriptional changes induced by *Vhl* and *Vhl/Pbrm1* mutations in proximal tubule cells are partly reflected in mouse and human ccRCC tumors

We next assessed the relevance of the transcriptional changes that are induced early after gene deletion to gene expression in established ccRCC tumors. We first compared the expression of genes that are up- or downregulated by *Vhl* deletion in male proximal tubule cells to differentially expressed genes in ccRCC tumors resulting from *Vhl/Trp53/Rb1*, *Vhl/Trp53/Rb1/Hif1a* or *Vhl/Trp53/Rb1/Hif2a* deletion in male mice (transcriptomes from female tumors are not available) compared to normal male renal cortex ^23^. Of the 300 genes that were upregulated by *Vhl* deletion, 142 were also upregulated in *Vhl/Trp53/Rb1* tumors (logfc > 1, *p* < 0.05 in both datasets), some of which were uniquely dependent on HIF-1α or on HIF-2α (Fig. 3e). Of the 97 genes that were downregulated by *Vhl* mutation, 44 were also downregulated in *Vhl/Trp53/Rb1* tumors (logfc < −1, *p* < 0.05 in both datasets) (Fig. 3e). We conclude that approximately 45% of the early transcriptomic changes induced by *Vhl* mutation are also present in *Vhl/Trp53/Rb1* mutant ccRCCs. We also took advantage of a new ccRCC tumor model (Witt *et al*., co-submitted manuscript) based on combined *Vhl/Trp53/Rb1/Pbrm1* mutation to assess whether the modifying effects of *Pbrm1* co-mutation on gene expression changes induced by *Vhl* mutation are also maintained in *Vhl/Trp53/Rb1* ccRCC tumors lacking *Pbrm1* compared to *Vhl/Trp53/Rb1* tumors with wild type *Pbrm1*. Only 16 of 186 genes (9%) that were upregulated and 49 of 310 genes (14%) that were downregulated upon *Pbrm1* loss in proximal tubule cells were also similarly dependent on *Pbrm1* in *Vhl/Trp53/Rb1* ccRCCs (logfc > 1 or < −1, *p* < 0.05 in both datasets) (Fig. 3f). This analysis suggests that at least some of the transcriptional changes that are induced early following *Pbrm1* mutation are maintained throughout tumor evolution.

To examine the possibility that there might be more extensive conserved gene expression changes of lower magnitude, we identified PBRM1-dependent, significantly upregulated (Fig. 3g) and downregulated (Fig. 3h) genes (FDR *q* < 0.05) without applying an expression fold cutoff in mouse proximal tubule cells, mouse ccRCC and human ccRCC (combined analysis of *PBRM1* wild type and mutant ccRCCs in the TCGA-KIRC and CPTAC datasets). This identified larger numbers of overlapping PBRM1-dependent differentially expressed genes between proximal tubules and either mouse ccRCC (132 up, 97 down) or human ccRCC (62 up, 90 down), from which 9 upregulated and 14 downregulated genes were present in all three settings. We conclude that at least some of the gene expression changes that are induced early after *Vhl/Pbrm1* mutation in mice are also present in established ccRCCs. Our analyses also suggest that some of the early gene expression changes may be lost during the process of tumor evolution, potentially due to different constellations of additional mutations that further modify the effects of loss of PBRM1 function on the transcriptome in each individual tumor.

### Many biological processes are differently impacted by *Vhl* and *Pbrm1* mutations and sex

To gain insight into how *Vhl* and *Vhl/Pbrm1* mutations may similarly or differently contribute to the earliest steps in ccRCC evolution in male and female mice, we conducted GSEA analyses of the proximal tubule RNA-seq datasets (Fig. 4a, Supplementary Data Table 4), as well as generated mRNA expression heatmaps of all differentially expressed genes that belong to these terms (Fig. 4b-j). As expected, the HALLMARK term Hypoxia was upregulated by *Vhl* and *Vhl/Pbrm1* mutation in male and female proximal tubule cells but at the individual gene level it was notable that some genes (*Siah2, F3, Selenbp2, Pdk3, Sap30, Csrp2, Gpc4, Ext1*) were only upregulated in male mice, and some *Vhl* loss-induced genes were dependent on *Pbrm1* only in male cells (*Mt1, Gpc3, Igfbp3, Pnrc1, Map3k1, Efna1*) (Fig. 4b). We probed the expression of glycolytic pathway genes which we have previously demonstrated are dependent on HIF-1α in *Vhl* mutant cells ^23,24^. All of these genes were upregulated in *Vhl* and *Vhl/Pbrm1* mutant male and female proximal tubule cells, with the exception of *Ndufa4l2*, which was upregulated in *Vhl* and *Vhl/Pbrm1* mutant male cells, but was only upregulated in *Vhl* mutant, but not in *Vhl/Pbrm1* mutant, female cells (Fig. 4c). NDUFA4L2 inhibits mitochondrial complex I and acts to reduce electron transport, oxygen consumption and the generation of reactive oxygen species ^41^. *Vhl/Pbrm1* mutant male and female proximal tubule cells are therefore predicted to differ in their capacity for mitochondrial oxidation, with male cells likely having a more strongly reduced capacity.

**Fig. 4.**
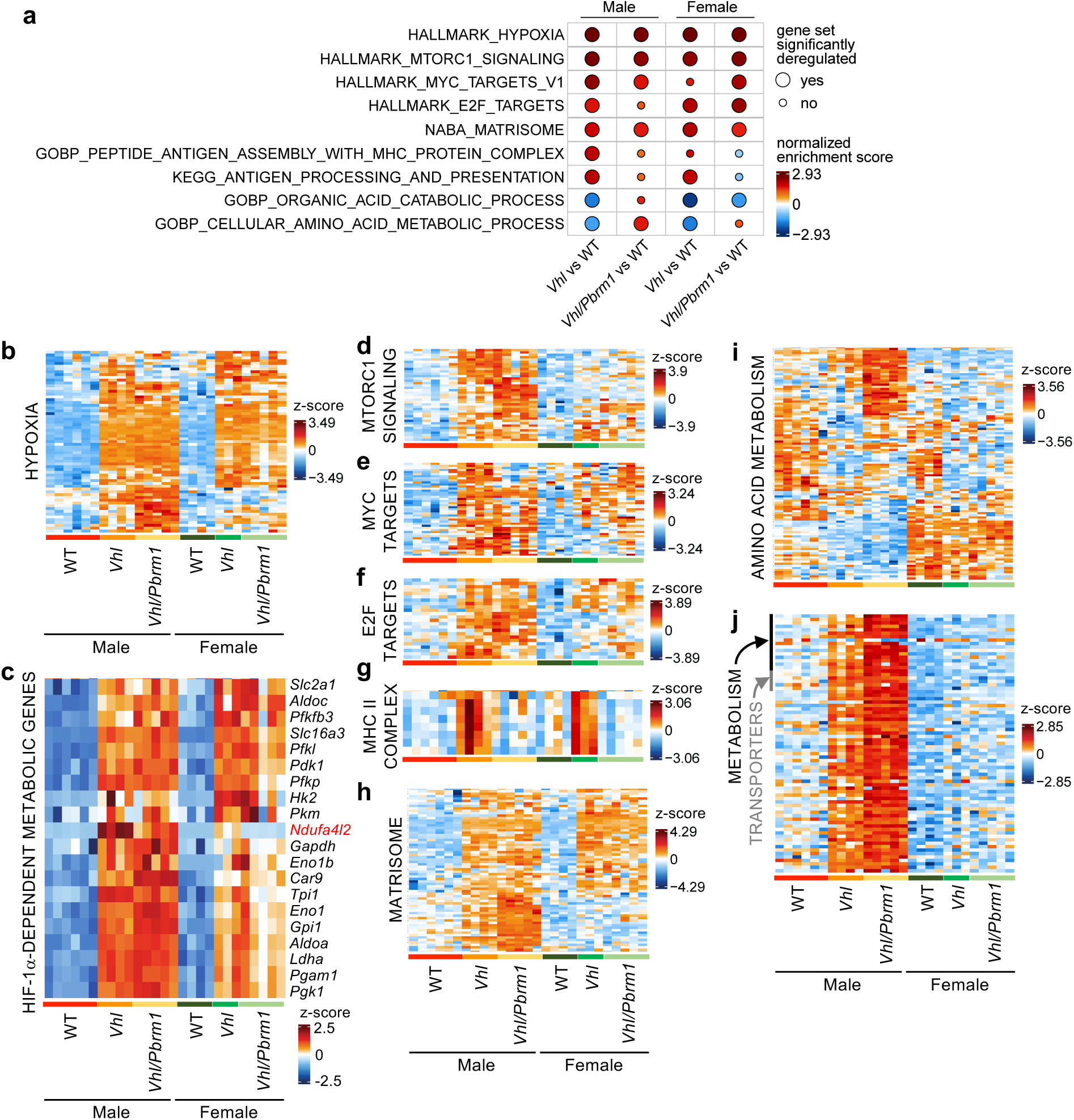
Sex-specific differential transcriptional effects of *Vhl* and *Vhl/Pbrm1* mutations affect multiple biological processes. **a** Selected differentially enriched GSEA terms in the comparisons of *Vhl* vs WT and *Vhl/Pbrm1* vs WT in male and female proximal tubule cells. **b-j** Gene expression heatmaps of differentially expressed genes that belong to the indicated gene sets in proximal tubules derived from independent mice (columns) of the indicated genotypes and sexes. For all heatmaps in b-j the gene lists and their expression values are provided in the Source Data Table.

Several, partly overlapping, HALLMARK terms that are associated with proliferation, including mTORC1 signalling, MYC targets v1 and E2F targets, were upregulated by *Vhl* and *Vhl/Pbrm1* mutation in both male and female proximal tubule cells, but in each of these terms there were several genes that were upregulated in males but not in females (Fig. 4d-f). These male-specific genes included *Gclc, Cyp51, Shmt2, Me1, Hmgcs1, Edem1* (for the mTORC1 signature), *Nap1l1, Gspt1, Hspd1, Etf1, Eif3j1, Hnrnpc, Psmc6, Eif2s1, Ctps, Cct4, Ifrd1* (for the MYC signature), and *Nap1l1, Gspt1, Ran, Pole4, Hus1, Ctps, Asf1a, Eif2s1* (for the E2F signature). A series of genes associated with antigen presentation via MHC class II (*Cd74, Ciita, H2-Aa, H2-Ab1, H2-DMa, H2-DMb1, H2-DMb2, H2-Eb1*) were upregulated in male and female *Vhl* mutant cells, but not in *Vhl/Pbrm1* mutant cells (Fig. 4g). The significance of this observation in the context of ccRCC development and shaping the anti-tumor immune response is investigated in a co-submitted manuscript (Witt *et al.*). The NABA term Matrisome was enriched by *Vhl* and *Vhl/Pbrm1* mutation in both male and female mice, however, there was a more complex pattern of expression at the level of individual genes (Fig. 4h). While some genes were upregulated in both males and females, others were upregulated by *Vhl* and *Vhl/Pbrm1* mutation only in male mice. These male-specific genes included those encoding extracellular signalling molecules (*Fgf1, Frzb, Il15, Il34, Ccl28, Pdgfd, Ntn1, Tnfsf10, Ptn*) and mediators of diverse other extracellular functions (*Ctsh, Creld2, Insl6, Gpc4, Mep1a, Mep1b, Vwa1, Col4a3, Col6a6, Tgm1, Hapln1*). Genes associated with the partly overlapping GOBP terms Organic Acid Catabolic Process and Cellular Amino Acid Metabolic Process showed differential GSEA enrichments between the *Vhl* and *Vhl/Pbrm1* mutant genotypes in both sexes. Particularly noteworthy are a set of 35 genes that were strongly upregulated only in *Vhl/Pbrm1* mutant male cells (Fig. 4i). Amongst these genes are several that participate in branched chain amino acid mitochondrial catabolism (*Bckdhb, Mccc1, Acad8, Echs1, Hibadh, Acat1*), tyrosine and tryptophan degradation (*Fah, Ido2, Aadat, Kyat1*) as well as glutathione and redox homeostasis (*Gclc, Gclm, Gss, Ggt1, Msra*), suggestive of a metabolic rewiring toward amino acid-driven mitochondrial energy generation and redox buffering, potentially to compensate for the reduced mitochondrial oxidation of glucose and fatty acids imposed by *Vhl* and *Vhl/Pbrm1* mutations in these cells. VHL-dependent changes in branched chain amino acid catabolism in human renal cancer cells have been reported ^42^. Finally, of the total set of genes that were specifically upregulated by *Vhl* or *Vhl/Pbrm1* mutation only in male cells, there was an enrichment for genes involved in diverse aspects of lipid, cholesterol and steroid metabolism (*Crot, Nudt19, Srd5a2, Hsd11b1, Hsd17b11, Acot12, Pm20d1, Ces2b, Cyp51, Agps, Lipo3*) as well as in uptake of omega-3 fatty acids (*Mfsd2a*), long chain fatty acids (*Cd36*), short chain fatty acids (*Slc5a8*), prostaglandins (*Slc22a7*), bile acids and steroids (*Slco1a4*) and riboflavins (*Slc52a3*) (Fig. 4j).

These analyses collectively show that additional *Pbrm1* mutation can modify the transcriptional changes induced by *Vhl* deletion, leading to either loss or gain of gene expression. Interestingly, these effects are also partly sex-dependent. The fact that these differently regulated genes participate in a diversity of biological pathways, including cellular metabolism, proliferation, antigen presentation and the matrisome, implies that the earliest steps of ccRCC development are likely to be different between males and females.

### *Vhl* and *Vhl*/*Pbrm1* mutations have sex-specific effects on lipid droplet accumulation

In addition to the above-described alterations in metabolic genes, *Vhl* and *Vhl/Pbrm1* deletions in both male and female cells caused upregulation of *Apoa4*, *Apoc3* and *Fabp5* (Fig. 5a), which are involved in intracellular lipid handling and transport, which we reasoned might either induce, or be a consequence of, altered lipid metabolism. We further reasoned that male *Vhl/Pbrm1* mutant proximal tubule cells might more strongly accumulate lipids than female cells due to the combination of i) their intrinsically higher natural high rate of mitochondrial activity, ii) induction of glycolysis, iii) the male-specific high expression of *Ndufa4l2* (Fig. 4c) which is predicted to reduce mitochondrial electron transport and iv) the male-specific upregulation of *Cd36*, *Mfsd2a* and *Slc5a8* (Fig. 5a), which are predicted to promote lipid and fatty acid uptake. These factors may cause a sustained imbalance in fatty acid uptake and mitochondrial oxidation, resulting in conversion into triacyl glycerol and storage in cytoplasmic lipid droplets. Histological comparisons of kidneys 2 weeks after the end of the period of tamoxifen feeding did not reveal any obvious lipid accumulation. Since we reasoned that it may take a longer time for sufficient lipids to accumulate, we generated cohorts at a timepoint of 20 weeks after feeding. Only male, but not female, *Vhl/Pbrm1* mutant tubules accumulated round, optically clear structures that are characteristic of lipid droplets (Fig. 5b). Lipid-droplet containing cells almost invariably stained positively for CFP/YFP (Fig. 5c) and we observed many examples of CFP/YFP-marked *Vhl/Pbrm1* mutant male, but not female, proximal tubule cells having lipid droplets while the adjacent wild type cells in the same tubule did not (Fig. 5d), demonstrating the cell autonomy of the phenotype. Immunohistochemical staining for perilipin 2 (PLIN2), which coats lipid droplets, revealed a strong accumulation of smaller lipid droplets only in *Vhl/Pbrm1* mutant, but not in *Vhl* mutant, tubules in males (Fig. 5e). The large optically-clear droplets did not stain positively for PLIN2, suggesting that these may represent a different cellular lipid pool. In females, PLIN2 accumulated in both *Vhl* and *Vhl/Pbrm1* mutant tubules (Fig. 5e), demonstrating a further sexual dimorphism in lipid handling induced by these tumor suppressor mutations.

**Fig. 5.**
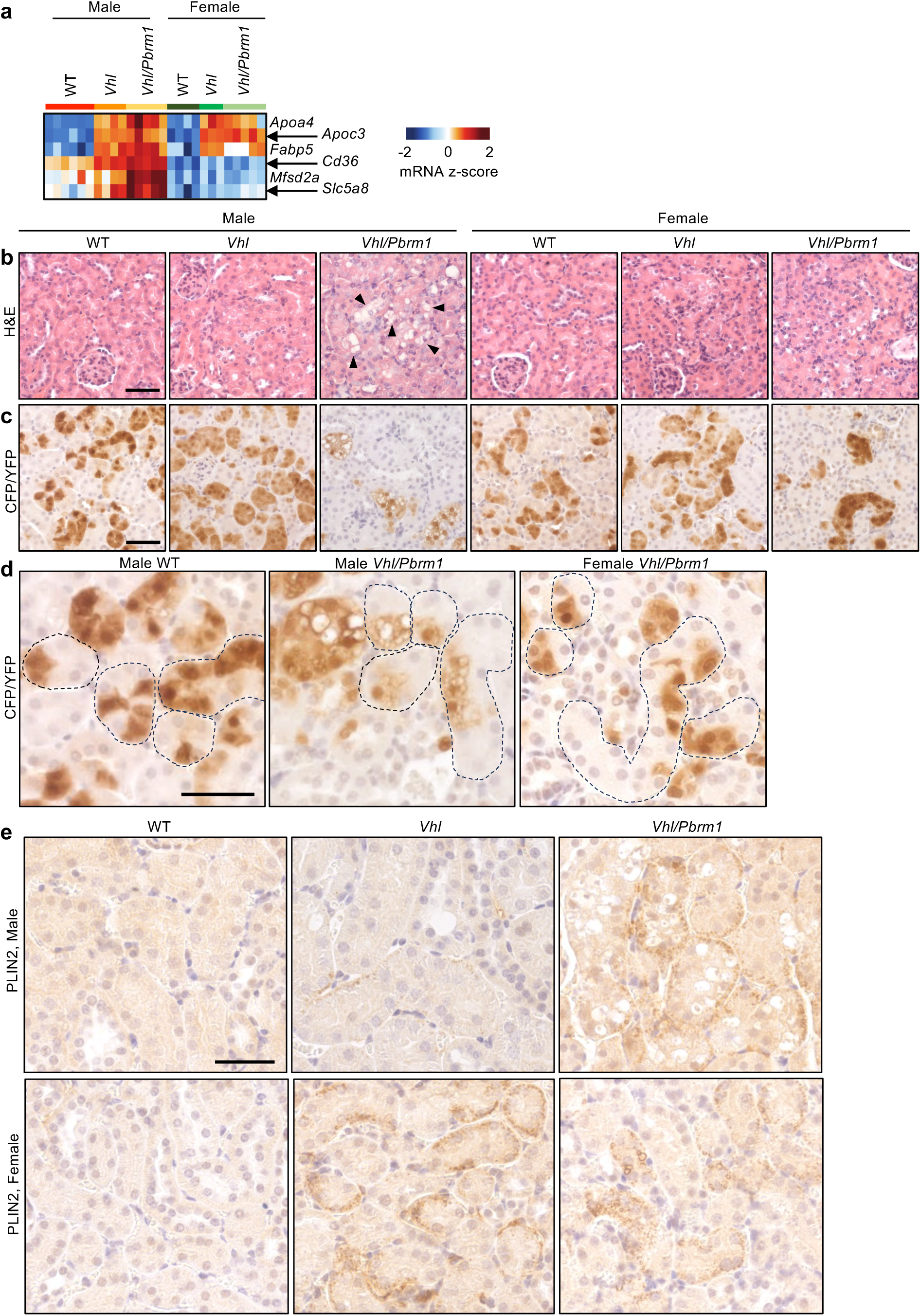
*Vhl* and *Vhl/Pbrm1* deletions have sex-specific effects on lipid accumulation in proximal tubule cells. **a** Gene expression heatmap of selected sex- or genotype-specific differentially expressed genes involved in lipid handling and uptake. **b** Representative H&E staining of kidneys from the indicated genotypes and sexes 20 weeks after tamoxifen feeding. Arrowheads indicate lipid droplets. **c** Representative immunohistochemical staining using an anti-CFP/YFP antibody to label cell clones that experienced Cre-mediated recombination. **d** High magnification zoom of individual WT male and *Vhl/Pbrm1* male and female kidney tubules (outlined by dotted lines) showing mosaic patterns of gene deletion as revealed by an anti-CFP/YFP antibody. **e** Immunohistochemical stains for PLIN2 in the indicated genotypes and sexes. Scale bars in b-e = 50 μm.

### *Vhl*/*Pbrm1* mutation enhances tubular repair, proliferation and clonal expansion in male proximal tubule cells

Two genes that are upregulated during the process of tubular repair, *Havcr1* (Fig. 6a), encoding Kidney Injury Molecule-1 (KIM-1) and *Fn1* (Fig. 6b), encoding fibronectin, were upregulated only in *Vhl/Pbrm1* male proximal tubules. Another tubular injury marker, *Ncam1* ^43,44^, encoding neural cell adhesion molecule 1, was upregulated by *Vhl* and *Vhl/Pbrm1* deletions in both male and female proximal tubules (Fig. 6c). Given these results, and since the accumulation of fatty acids and lipids can induce lipotoxic cell death of proximal tubule cells, which can trigger the process of compensatory tubular repair ^45,46^, we quantified the protein expression of KIM-1 and the SOX9 transcription factor, which serve as molecular markers of ongoing cellular stress and tubular repair ^47,48^. In males, the number of tubules with at least one KIM-1 positive cell (Fig. 6d) and the frequency of cells expressing SOX9 (Fig. 6e) were elevated only in *Vhl/Pbrm1* mutants, consistent with the accumulation of PLIN-2-coated and larger lipid droplets in these cells and with the idea of lipotoxicity-induced tubular damage. In females, *Vhl* deletion alone increased the frequency of KIM-1 and SOX9 positivity and positivity for these markers was further increased in *Vhl/Pbrm1* mutant cells (Fig. 6d,e). These results correlate with the accumulation of PLIN-2-coated lipid droplets in these genotypes. Also consistent with both of these sets of results, staining for the proliferation marker Ki-67 revealed that only *Vhl/Pbrm1* mutation, but not *Vhl* mutation, induced ongoing proliferation in males, while both *Vhl* mutation and *Vhl/Pbrm1* mutation caused elevated proliferation of proximal tubule cells in females (Fig. 6f). In conclusion, there is a correlation between metabolic gene expression changes, accumulation of PLIN2-positive lipid droplets, tubular repair and ongoing proliferation in both male and female cells, but the gene mutational dependency of these phenotypes differs between the sexes. While either *Vhl* or *Vhl/Pbrm1* mutation induced these responses in female mice, in males, only *Vhl/Pbrm1* mutation increased PLIN2 positivity, tubular repair and resulted in the formation of large lipid droplets.

**Fig. 6.**
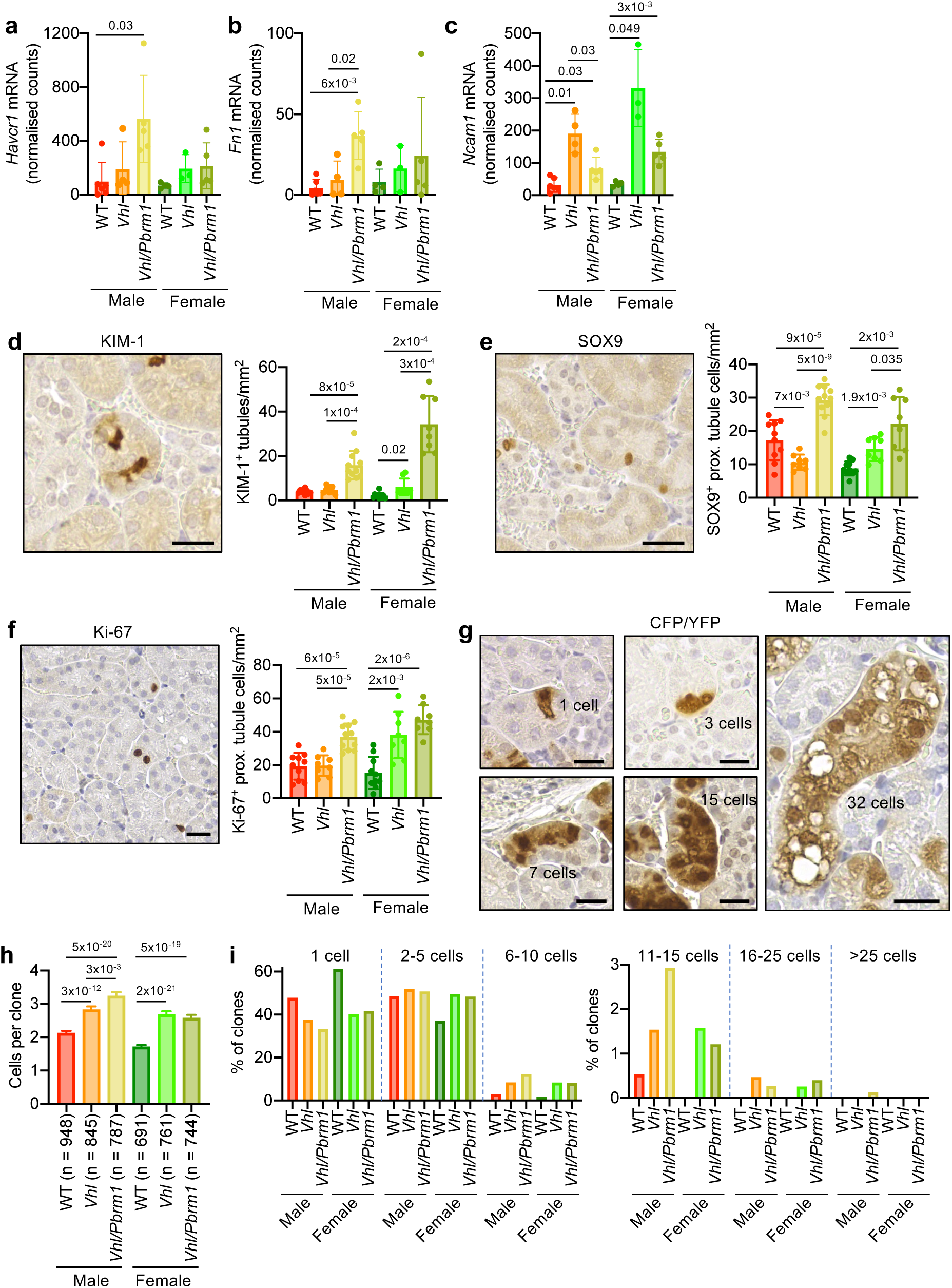
*Vhl* and *Vhl/Pbrm1* deletions induce tubular repair, proliferation and clonal expansion. **a-c** Relative mRNA abundance (normalised reads from RNA-seq) of *Havcr1* (**a**), *Fn1* (**b**) and *Ncam1* (**c**) in proximal tubule cells of the indicated genotypes and sexes. Data represents mean ± std. dev. of 4-6 mice. *P* values were calculated by two-sided unpaired t-test with Welch’s correction. **d** Representative immunohistochemical staining for KIM-1 20 weeks after tamoxifen feeding and quantification of the number of tubules with one or more KIM-1 positive cells per mm^2^. **e** Representative immunohistochemical staining for SOX9 20 weeks after tamoxifen feeding and quantification of the number of SOX9 positive proximal tubule cells per mm^2^. **f** Representative immunohistochemical staining for Ki-67 20 weeks after tamoxifen feeding and quantification of the number of Ki-67 positive proximal tubule cells per mm^2^. Data in d-f represents mean ± std. dev. derived from analyses of 8-11 kidneys of each genotype and sex. *P* values were calculated by two-sided unpaired t-test with Welch’s correction. **g** Examples of CFP/YFP positive proximal tubule clones of different sizes 20 weeks after tamoxifen feeding. Scale bars in d-g = 20 μm. **h** Mean ± SEM clone sizes 20 weeks after tamoxifen feeding. 6-8 kidneys of each genotype and sex were analysed and the total number of clones analysed is shown. **i** Distributions of proximal tubule clone sizes in each genotype and sex.

Tubular repair and proliferation could be one driving force that contributes to clonal expansion of mutant cells at the earliest stages of cancer evolution. To test this idea, we took advantage of the fact that Cre-mediated recombination of the *Ubow* transgene marks the initial cell in which recombination took place as well as all of the potential daughter cells that arise over time, allowing us to compare the size distributions (Fig. 6g) of clones 20 weeks after tamoxifen feeding across the different genotypes and sexes. Compared to wild type mice, *Vhl* mutation increased the average proximal tubule clone size by 33% in males and 57% in females (Fig. 6h), consistent with the previously described early, but not sustained, proliferative effect of *Vhl* deletion ^40,49^. *Vhl/Pbrm1* deletion further increased the average clone size in males (52% higher than in wild type) but not in females (51% higher than in wild type) (Fig. 6h). These results are concordant with the Ki-67 analyses which provide a snapshot of proliferation; *Vhl/Pbrm1* male mice show elevated levels of proliferation and increased clone size compared to *Vhl* mutation, while both *Vhl* and *Vhl/Pbrm1* female mice show similarly elevated levels of proliferation and similar increases in clone size. These data are reflected in the distributions of clone sizes (Fig. 6i), with both *Vhl* and *Vhl/Pbrm1* deletions in both male and female mice causing a reduction in the number of single-cell clones, together with increases in the frequencies of clones with 2-5 and 6-10 cells. *Vhl/Pbrm1* mutation exhibited a differential effect in increasing the frequency of clones with 6-10 and with 11-15 cells only in male, but not female, mice. Although present at very low frequency, *Vhl/Pbrm1* mutant males were also the only condition where clone sizes above 25 cells were observed. These data are consistent with additional *Pbrm1* mutation promoting the initial *Vhl* loss-induced expansion of mutant cells preferentially in male mice, particularly in terms of causing a relatively rare but more robust clonal expansion. These data illustrate that there are sexually dimorphic effects of *Vhl/Pbrm1* mutation on clonal expansion, a very early step in the process of tumor evolution.

## DISCUSSION

This study identified that the starting points of ccRCC formation in male and female mice are different. Proximal tubule cells display intrinsic sex-specific gene expression patterns, particularly in genes that control cellular metabolism, mitochondrial abundance and protein translation. We further identified that male and female proximal tubule cells responded differently to mutation of ccRCC truncal driver genes. There was only a partial overlap of the transcriptional effects of *Vhl* or *Vhl/Pbrm1* mutation in male and female cells. *Pbrm1* mutation modulated the transcriptional response to *Vhl* deletion more strongly in male than female cells, and resulted in either loss or gain of gene expression. Collectively, the intrinsically sex-specific genes and the sex-specific, mutation-induced gene expression patterns are predicted to affect cellular metabolism, mitochondrial activity, cellular proliferation, protein translation, antigen presentation and a diversity of extracellular processes, implying that the earliest steps of tumor development might be substantially different between males and females.

We identified several, potentially interrelated, sex-specific consequences of *Vhl/Pbrm1* mutation. While *Vhl* and *Vhl/Pbrm1* mutation in female proximal tubule cells induced the accumulation of PLIN2-coated lipid droplets, this phenotype was observed only in *Vhl/Pbrm1* mutant, but not *Vhl* mutant, proximal tubule cells in male mice. Gene expression patterns also predicted that *Vhl/Pbrm1* mutant male proximal tubules might exhibit a stronger imbalance in the import and mitochondrial oxidation of fatty acids, resulting from the combination of high male-specific intrinsic mitochondrial abundance and substrate oxidation activity, the HIF-1α-dependent induction of a switch to glycolytic metabolism, the inhibition of mitochondrial electron transport complex I via *Ndufa4l2* upregulation and the *Vhl/Pbrm1*-dependent upregulation of genes that are predicted to promote lipid and fatty acid uptake. Indeed, only male *Vhl/Pbrm1* mutant proximal tubule epithelial cells accumulated large, optically-clear lipid droplets, reflecting the hallmark clear cell histological feature of this tumor type. It is known that accumulation of lipid droplets can induce lipotoxicity and death of proximal tubule cells, which can trigger the process of compensatory tubular repair ^45,46^. Consistent with this idea, both *Vhl* and *Vhl/Pbrm1* mutant proximal tubules in females, but only *Vhl/Pbrm1* mutant proximal tubules in males, showed elevated frequencies of cells expressing markers of ongoing tubular repair and proliferation. We speculate that this process could represent one factor that might contribute to the initial expansion of mutant cells. An enhanced propensity of lipid-laden mutant cells to undergo lipotoxic cell death might cause an increase in the frequency of compensatory division of neighboring mutant cells, resulting in increased clone sizes over time. It is noteworthy that while both male and female tubular cells exhibited moderate clonal expansion in response to *Vhl* mutation alone, which might potentially relate to the observed upregulation of pro-proliferative mTORC1, MYC and E2F gene expression signatures, additional *Pbrm1* mutation caused a further increase in clone size only in males and not in females. We note that the increase in clonal expansion induced by additional *Pbrm1* mutation in males is not a very large effect (an additional 20% increase in average clone sizes over *Vhl* mutant cells), however, over time, this small increase in cell proliferation is expected to result in an expanded pool of mutant cells, increasing the chances of the accumulation of additional cooperating mutations that then further drive the process of tumor evolution. This is consistent with the long timeframe of tumor evolution in this mouse model as well as in human ccRCC ^37^. While there were no sex-specific differences in the low frequency of ccRCC development observed in our model of *Vhl/Pbrm1* deletion, in a co-submitted manuscript (Witt *et al.*) we demonstrate that *Pbrm1* mutation in the genetic background of the *Vhl/Trp53/Rb1* mutant model of ccRCC moderately accelerates tumor onset in male but not in female mice, supporting the notion that *Pbrm1* mutation may differently affect tumor formation in males and females. The fact that *PBRM1* mutations are more prevalent in ccRCCs from men than from women ^51^ is consistent with a more potent ccRCC tumor-promoting effect of *PBRM1* mutation in male proximal tubule cells.

In this study we chose to employ bulk RNA-seq of sorted cells rather than single cell RNA-seq to take advantage of higher sequencing depth to more reliably detect changes in the expression of lowly expressed genes, which we deemed particularly important for identifying downregulated genes that we anticipated might result from *Pbrm1* mutation. A limitation of this approach is that it does not provide the resolution of the effects of gene mutations on different sub-types of proximal tubule epithelial cells, which have been previously demonstrated for *Vhl* mutation ^40,49^. Our findings and mouse models nonetheless set the stage for future experiments that could take advantage of single cell-resolution spatial transcriptomics to assess whether mRNA expression patterns might change throughout different stages of cancer evolution. Our transcriptomic data predicts that male and female *Vhl* and *Vhl/Pbrm1* mutant cells have specific metabolic profiles. Beyond the accumulation of lipid droplets, we were not able to prove that these metabolic changes take place *in vivo* for several technical reasons. Limitations of the yield of viable sorted cells prevented metabolomic analyses and spatial metabolomics approaches do not offer the necessary single-cell resolution. We investigated whether Adeno-Cre-mediated deletion of *Vhl/Pbrm1* in cultured primary renal epithelial cells could represent a cell culture model to allow the study of metabolic changes. However, *Vhl/Pbrm1* deletion from cultured male and female renal epithelial cells showed a very low degree of overlap in terms of differentially expressed genes (Supplementary Fig. 5a,b) and of overall transcriptional profiles (Supplementary Fig. 5c,d) with the directly isolated cells. While this highlights the relevance of our analyses of genetically-labelled cells isolated directly from kidneys, it also demonstrates that culture systems do not adequately reflect the *in vivo* setting in a way that allows the accurate study of processes such as cellular metabolism that are relevant for tumor formation. Finally, while our data demonstrate a correlation between lipid accumulation, tubular repair, proliferation and clonal expansion, they do not establish causality and do not exclude that other processes such as alterations in cell signaling, cell cycle and cell death may also contribute to the differing effects of *Vhl* and *Vhl/Pbrm1* mutations on early tumor formation in male and female cells. These processes may act separately to, or in concert with, other sex-specific biological factors such as hormones or genes on the sex chromosomes that likely also play a role in tumor incidence disparities ^50^.

Overall, our findings highlight how genetic mutations can have sex-specific molecular and cellular effects that are relevant for cancer biology. We show that there are important molecular differences between the male and female cells of origin of ccRCC in their response to mutational inactivation of the mouse homologues of the two most commonly mutated human ccRCC driver genes. It is therefore conceivable that similar sexual dimorphisms in the earliest steps of ccRCC evolution exist in humans and that these could contribute to the sex-bias in cancer incidence. Most tumor types at shared anatomical sites arise more frequently in men than in women, and this cannot be explained solely by sex-specific differences in behavioral or environmental risk factors ^2^. We suggest that the possibility that there are similar sex-specific differences in the responses of cancer cells of origin to cancer-specific driver mutations warrants more intensive investigation across a wide range of tumors.

## METHODS

### Mice

Previously-described *Ksp1.3-CreER^T^*^2^ (C57BL/6J background) ^33^, *Vhl^fl/fl^* (C57BL/6J background) ^52^, *Pbrm1^fl/fl^* (B6;129 background) ^53^ were intercrossed to generate the experimental mouse lines for the long term studies and for proximal tubule epithelial cell cultures. These lines were crossed with *Ubow* (C57BL/6J background) ^38^ mice to generate the strains for the clonal tracking experiments. All final *Ubow* strains contained > 98.5% C57BL/6J genetic background. Littermate mice that lacked the Cre transgene served as wild type controls. Gene deletion in 6 week-old mice was achieved by feeding *ad libitum* with tamoxifen-containing food A155T70401 (400 parts per million, Ssniff) for 2 weeks. Mice were fed thereafter with normal diet D12450Hi (10 kcal% fat, 70 kcal% carbohydrate, 20 kcal% protein, Research Diets Inc). All animal experimental protocols were approved under the licenses G-19/16 and G-21/091 of the Regierungspräsidium Freiburg and all methods were carried out in accordance to these regulations. Investigators were not blinded to the genotype of the mice.

### Immunohistochemistry and immunofluorescence

Mouse kidneys were fixed in 10% formalin, paraffin-embedded and cut in 5 µm sections. Immunofluorescence and immunohistochemical stainings were performed as previously described ^54^. Primary antibodies against the following proteins or epitopes were used at the following dilutions and antigen retrieval conditions: CA9 (1:150, Invitrogen, PA1-16592, citrate, 15 min, 114°C), GFP (1:2000 for IHC, 1:200 for IF, Abcam, ab5450, citrate, 15 min, 114°C), Ki-67 (1:1000, Abcam, ab15580, citrate, 15 min, 114°C), PLIN2 (1:400, Proteintech, 15294-1-AP, citrate, 15 min, 114°C), HAVCR1/KIM-1 (1:300, R&D Systems, AF1817, EDTA/0.05% Tween pH 9.0, 15 min, 114°C), SOX9 (1:1000, Abcam, ab185230, EDTA/0.05% Tween pH 9.0, 15 min, 114°C), LTL-FITC (1:250, Vector Laboratories, VEC-FL-1321). For quantification, stained tissue sections were scanned with the Zeiss Axio Scan.Z1 slide scanner at a magnification of 20x (NA 0.50) and images were imported into the QuPath software for further analysis. The area of the kidney cortex was determined by manually contouring the borders and the number of positive cells or tubules was counted manually.

### Multiplex OPAL immunofluorescence

Kidney sections were de-paraffinized and processed for multiplex staining as follows: antigen retrieval was performed in a pressure cooker using EDTA/0.05% Tween (pH 9.0) as antigen retrieval buffer (15 minutes at 114°C). Two blocking steps were performed, 15 minutes in 3% HOand 20 minutes in blocking solution (included in Opal 3-Plex Anti-Rb Manual Detection Kit, Akoya/Quanterix, NEL840001KT) respectively. All primary antibodies were diluted in blocking solution (Akoya/Quanterix) and incubated either for 1h at room temperature (RT) or over-night at 4°C. Primary antibodies used: anti-GFP (1:1000, Abcam, ab5450), anti-MEGALIN/LRP2 (1:500, Abcam, ab76969), anti-PBRM1 (1:1000, Bethyl, A301-591A). Primary antibody incubation was followed by secondary anti-rabbit HRP (1:1000, Invitrogen, #31460,) incubation for 20 minutes at RT, followed by OPAL dye (520, 570, 620 or 690, diluted in Akoya amplification diluent) incubation for 10 minutes at RT. For each consecutive antibody the protocol was repeated from antigen retrieval through to OPAL dye incubation. After staining completion, the sections were counterstained against DAPI (Akoya/Quanterix) and mounted using Fluoromount G (Invitrogen). Slides were scanned using Akoya Phenocycler Fusion and images processed using the InForm software and ImageJ.

### Proximal tubule cell isolation by flow cytometry

Primary kidney epithelial cells (PKCs) were isolated from male and female mice that were fed tamoxifen containing food for 2 weeks, followed by additional 2 weeks on normal chow. Five to seven mice per genotype were sacrificed and their kidneys were isolated. After removing the capsule, dissected kidneys were minced with a scalpel and the cell mixture was transferred to a gentleMACS^TM^ C-tube (Miltenyi Biotec). Multi Tissue Dissociation Kit 2 (Miltenyi Biotec) was used in the following volumes: 4.8 ml of Buffer X, 50 μl of enzyme P, 50 μl of buffer Y, 100 μl of enzyme D and 20 μl of enzyme A per 1 g of tissue were mixed and added to the kidney tissue. The tissues were processed with the preloaded program m_spleen_04, followed by m_brain_03 and finally program 37_MultiE of the gentleMACS^TM^ Dissociator. The digested and homogenized kidney solution was filtered through a 70 µm cell strainer, washed with 15 ml of PBS and after centrifugation lysis of red blood cells with 1 ml of ACK buffer (Thermo Fisher) was performed for 1 minute at RT. ACK buffer was inactivated with 10 ml of HBSS supplemented with 5% FCS. After centrifugation the cell pellets were resuspended with primary antibody solution: biotinylated LTL (1:500, Vector Laboratories, B-1325) diluted in MACS buffer (PBS supplemented with 372 mg EDTA and 2% FCS) and incubated for 30 minutes on ice. Thereafter, samples were washed with MACS buffer and centrifuged. Pellets were resuspended in secondary antibody solution: Streptavidin-Alexa Fluor^TM^ 700 conjugate (1:1,000, Thermo Scientific, S21383) in MACS buffer and incubated for 30 minutes on ice in the dark. After subsequent washing and centrifugation, the pellets were resuspended in an appropriate volume of MACS buffer for cell sorting. Cell sorting was performed on a BC CytoFlex SRT with a nozzle size of 100 µm. Double positive LTL (LTL^+^) and CFP/YFP (CFP^+^/YFP^+^) cells were collected (600-12,000 cells per mouse), as well as LTL negative cells (LTL^−^) (50,000 cells per mouse). Cell pellets were frozen in 75 µl Buffer RLT (RNeasy Micro Kit, Qiagen) supplemented with 1:100 β-mercaptoethanol and stored for up to two weeks at −80°C before RNA isolation was performed.

### Primary renal epithelial cell culture

Primary renal epithelial cells were isolated from 9-12 week-old male and female *Vhl^fl/fl^Pbrm1^fl/fl^* mouse kidneys using the protocol described above, with the only difference that the tissues were processed only with the preloaded program 37_MultiE of the gentleMACS^TM^ Dissociator. Following ACK lysis, cell pellets were resuspended in 20 ml K-1 medium (Dulbecco’s modified Eagle’s medium) and Hams F12 mixed 1:1, 2 mM glutamine, 10 kU/ml penicillin, 10 mg/ml streptomycin, hormone mix (5 μg/ml insulin, 1.25 ng/ml prostaglandin E1 (PGE1), 34 pg/ml triiodothyronine (T3), 5 μg/ml transferrin, 1.73 ng/ml sodium selenite, 18 ng/ml of hydrocortisone, and 25 ng/ml epidermal growth factor) + 0.5 % FCS. Cells were counted and seeded at 5×10^5^ cells per 6 cm dish. Cells were cultured in a humidified 5% (v/v) COand 20% Oincubator at 37 °C. Medium was changed 24**-**48 h after plating and cells were infected with Adenoviruses Type 5 (dE1/E3), expressing the Cre recombinase + GFP (Adeno-Cre-GFP, Vector Biolabs, 1700) or GFP only (Adeno-GFP, Vector Biolabs, 1060) 3 to 4 days after isolation (at 30% cell confluency approximately). Cells were never passaged and were directly harvested for protein or RNA 4 days post infection with Adenovirus.

### RNA sequencing and data analysis

RNA purification and preparation for RNA sequencing was performed according to the manufacturer’s instructions of the RNeasy Micro Kit (Qiagen) for the sorted primary kidney cells or NucleoSpin RNA kit (Macherey Nagel) for the cultured primary kidney cells. For the sorted primary kidney cells SMART-Seq cDNA synthesis followed by Nextera library preparation and sequencing (paired-end, 150bp, Illumina NovaSeq X Plus instrument) were performed by Novogene. For cultured primary kidney cells, library preparation (Illumina TruSeq Stranded RNA Library Preparation kit) and sequencing (paired-end, 100bp for male and 150bp for female samples, Illumina NovaSeq 6000 instrument) were performed by the core facility of the DKFZ (German Cancer Research Center) in Heidelberg. Quality of sequencing reads was assessed with FastQC (v0.11.9, http://www.bioinformatics.babraham.ac.uk/projects/fastqc/). Raw sequencing reads were processed with cutadapt (v4.2) ^55^ to trim low-quality bases, trim adapter sequences as well as poly-G artifacts, remove low-quality reads (more than 2 expected errors and/or more than 30% N-bases) and remove short reads (less than 20 nucleotides). Processed reads were aligned against the mouse reference genome GRCm39 from Ensembl (v108) ^56^ using STAR (v2.6.1e) ^57^. The number of reads per gene was counted during the alignment process, using the STAR option “--quantMode Gene Counts”. Subsequent data analysis was executed with R (v4.2.1). Samples of WT *Ubow* mice, *Ksp1.3-CreER^T^*^2^ *Ubow* mice and cultured primary renal epithelial cells were analyzed separately. Gene count normalization and differential gene expression analysis were performed with the R packages limma (v3.52.4) ^58^ and edgeR (v3.38.4) ^59^. Genes were considered differentially expressed with *p*_adj_< 0.05. The calculated t-statistic values were used as input values for the gene set enrichment analysis. Gene set enrichment analysis was carried out with the R function “GSEA” of package clusterProfiler (v4.4.4) ^60^, based on a selection of gene sets retrieved with the R package msigdbr (v7.5.1) (included gene set collections: hallmark, canonical pathways excluding WikiPathways, transcription factor targets, Gene Ontology). Gene sets were considered differentially regulated with *p*_adj_< 0.05. Raw RNA-seq data has been uploaded to GEO with identifier GSE330840 (https://www.ncbi.nlm.nih.gov/geo/query/acc.cgi?acc=GSE330840, reviewer token olutkcuijjcdzcj).

### Assessment of knock-out efficiency in RNA sequencing samples

The number of reads aligned to a single exon (or the first intron of *Vhl*) of *Vhl* or *Pbrm1* was determined with samtools coverage (v1.14) ^61^ using the mouse reference genome GRCm39 annotations of Ensembl (v108) ^56^. The counts of all unfloxed exons of *Vhl* or *Pbrm1* were summed up. For *Vhl*, this yielded one floxed exon count (exon 1), one intron count (intron 1), and one unfloxed count per sample. For *Pbrm1*, this yielded one floxed exon count (exon 18) and one unfloxed count per sample. Count values were library size-adjusted with the size factors calculated during differential gene expression analysis, and logCPM-transformed.

### CellMatchR analyses

In order to verify the cellular identity of sorted proximal tubule cells we used the online tool CellMatchR ^39^ (https://nephgen2024.shinyapps.io/CellMatchR/). CellMatchR compares the genetic expression profile obtained from bulk RNA-sequencing from cell lines, primary cells or whole tissue to kidney single cell RNA-sequencing references. Bulk RNA-sequencing data (normalized counts per million) was cross-compared to renal cell type specific gene lists using a rank-based Spearman’s correlation between the samples and reference samples. The most similar cell types are displayed at the top of the list.

### Western blotting

Cultured cells were lysed in RIPA (Radioimmunoprecipitation assay) buffer (50 mM Tris–HCl at pH 8.0, 150 mM sodium chloride, 1% (v/v) NP-40, 0.5% (w/v) sodium deoxycholate, 0.1% (w/v) SDS (sodium dodecyl sulfate), 5 mM sodium fluoride, 1 mM sodium orthovanadate, 1mM PMSF (phenylmethylsulphonyl fluoride) and Protease Inhibitor Cocktail (1:100, Sigma Aldrich). Western blots were probed with the following antibodies anti-pVHL(m)(1:2000, ref. ^62^), anti-PBRM1 (1:2000, Bethyl Laboratories, A301-591A), anti-β-ACTIN (1:5000, Sigma Aldrich, A2228), anti-VINCULIN (EPR8185) (1:10000, Abcam, ab129002).

### Comparison of mouse and human RNA-seq data

RNA-seq counts and somatic WES variant calls for TCGA-KIRC and CPTAC ccRCC were obtained from the Genomic Data Commons (https://portal.gdc.cancer.gov/). Counts were filtered to protein-coding genes (minimum 10 counts in ≥15% of samples) and normalized using TMM-scaled logCPM (edgeR). Analysis was restricted to *VHL*-mutated samples (TCGA n = 150, CPTAC n = 130). Differential expression between *PBRM1*-mutant and wild-type samples was tested per cohort using the Wilcoxon rank-sum test and combined via sample-size-weighted Fisher’s method across 16,143 shared genes. Combined *p*-values were adjusted for multiple testing using the Benjamini-Hochberg method. Mouse-to-human ortholog mapping was performed using one-to-one orthologs from the HomoloGene database (NCBI).

### Statistics

The following statistical tests were employed in this study: Gene expression values and immunohistochemical quantifications were tested for normality and differences between pairs were determined using two-sided Student’s t-test without adjustment for multiple comparisons, rank-based Spearman’s correlation was used to assess similarity of gene expression of sorted samples to cell-type specific gene expression, gene count normalization and differential gene expression analysis were performed with the R packages limma (v3.52.4) and edgeR (v3.38.4), gene set enrichment analysis was carried out with the R function “GSEA” of package clusterProfiler (v4.4.4) and Wilcoxon rank-sum test adjusted for multiple testing using the Benjamini-Hochberg method was used to investigate differential gene expression in human ccRCC. *P* < 0.05 or *P*< 0.05 was considered statistically significant.

## DATA AVAILABILITY

Source data are provided with this paper. Unprocessed RNA sequencing data have been uploaded to GEO with identifier GSE330840. Normalised reads of the RNA-seq dataset are provided in Supplementary Data Tables 1 and 2, GSEA results are provided in Supplementary Data Tables 3 and 4. The source data underlying all other heatmaps and graphs are provided in the Source Data file. All remaining relevant data are available in the article, supplementary information, or from the corresponding author upon reasonable request.

## ACKNOWLEDGEMENTS

This work was funded by grants from the Deutsche Forschungsgemeinschaft (DFG, German Research Foundation) SFB1479 – Project ID 441891347 (to I.J.F, M.B., A.K.), SFB1453 - Project ID 431984000 (to I.J.F, M.B., P.S., A.K.). SFB1160 – Project ID 256073931 (to M.B.), TRR167 – Project ID 259373024 (to M.B.), TRR353 – Project ID 471011418 (to M.B.), TRR417 – Project ID 540805631 (to M.B.), and FOR5476 – Project-ID 493802833 (to I.J.F., M.B.). This work was funded by the German Federal Ministry of Education and Research (BMBF) PM4Onco–FKZ 01ZZ2322A (to M.B., P.M.). The work of P.S. is supported by the DFG Project-ID 530592017 (SCHL 2292/3–1) and P.S. and A.K. are supported by Germany‘s Excellence Strategy (CIBSS – EXC-2189 – Project-ID 390939984). We are grateful to the staff of the Lighthouse Core Facility for assistance with cell sorting and microscopy (funded in part by the Medical Faculty, University of Freiburg (2021/B3-Fol) and the DFG (Project ID 450392965) and are grateful to the Core Facility AMIR (DFG-RIsources N° RI_00052) for support with animal imaging. The Proteomic Platform – Core Facility (ProtCF) was supported by the Medical Faculty of the University of Freiburg to Prof. Dr. Oliver Schilling (2021/A3-Sch and 2023/A3-Sch).

## AUTHOR CONTRIBUTIONS

Conceptualisation, A.C., C.K., K.M., S.J., I.J.F., M.A.,

Methodology, Formal Analysis and Investigation, A.C., C.K., K.M., S.J., M.M., Y.F., C.W., F.C., A.W., M.S., M.A.

Writing, I.J.F., M.A., A.C., C.K.,

Visualization, A.C., C.K., K.M., C.W., F.C., A.W., I.J.F., M.A.,

Supervision, M.A., I.J.F., M.B., P.S., S.H., A.K.,

Funding Acquisition, I.J.F., M.B., P.S., A.K.

---

**Supplementary Fig. 1.**
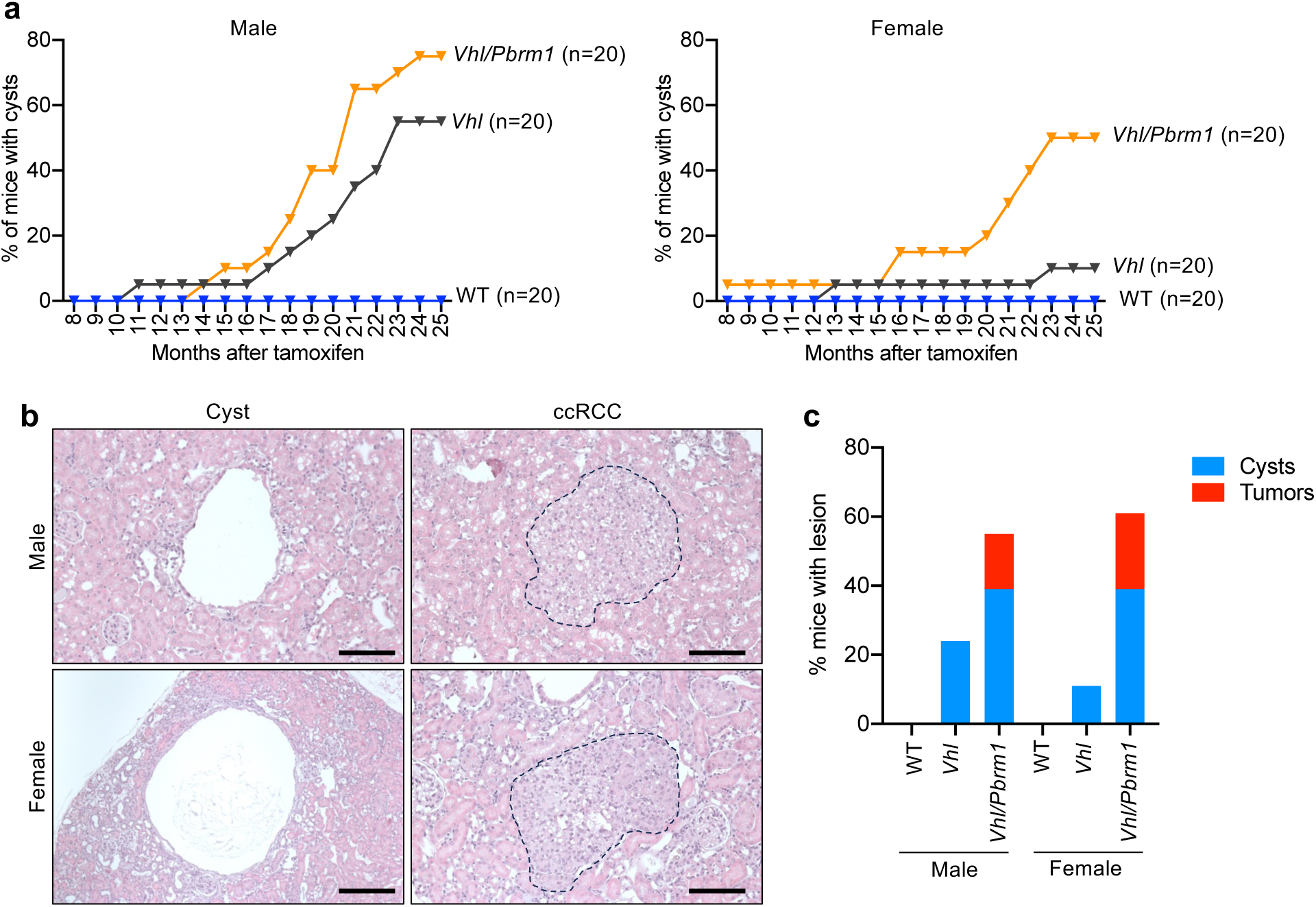
*Vhl/Pbrm1* deletion causes ccRCC evolution. **a** Percentage of mice of each genotype and sex that developed cysts visible by longitudinal ultrasound imaging. **b** Examples of cysts and ccRCCs that arose in male and female *Vhl/Pbrm1* mutant mice. Scale bars = 100 μm. **c** Quantifications of the percentage of mice of the indicated genotypes and sexes that exhibited cysts or ccRCCs in histological sections of kidneys 25 months after tamoxifen feeding.

**Supplementary Fig. 2.**
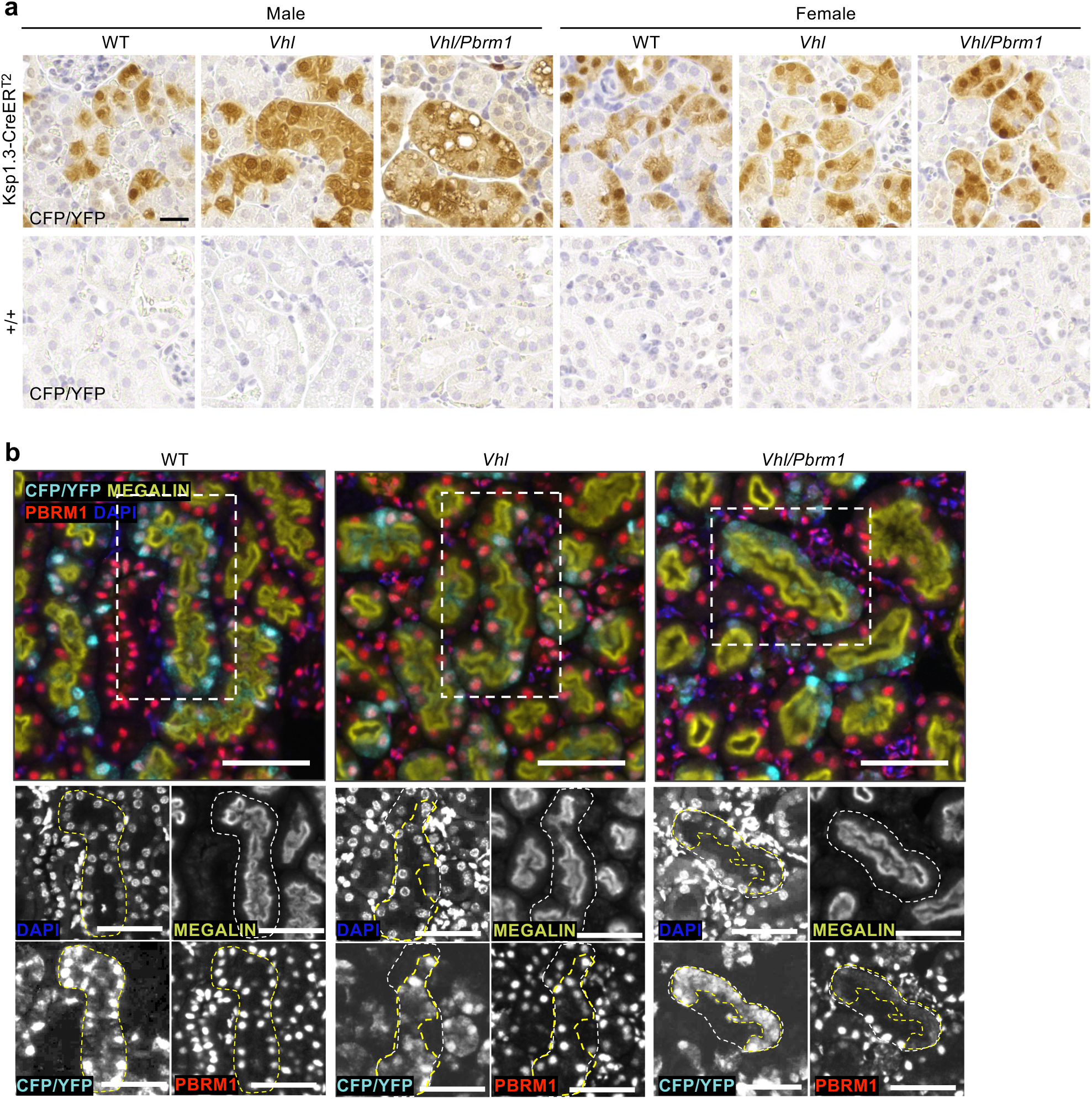
Validation of the specificity of clone detection and split channel images related to Fig. 1e. **a** Representative anti-GFP immunohistochemical stainings in tamoxifen-fed Cre-expressing (Ksp1.3-CreER^T2^) and Cre-negative (+/+) pairs of littermate mice of the indicated genotypes. Scale bar = 20 μm. **b** Extended data related to Fig. 1e. Black and white images depict the signals from the different channels in the regions marked with a dotted box. Individual tubules are outlined with white dotted lines and clones are outlined with yellow dotted lines. Scale bar = 50 μm.

**Supplementary Fig. 3.**
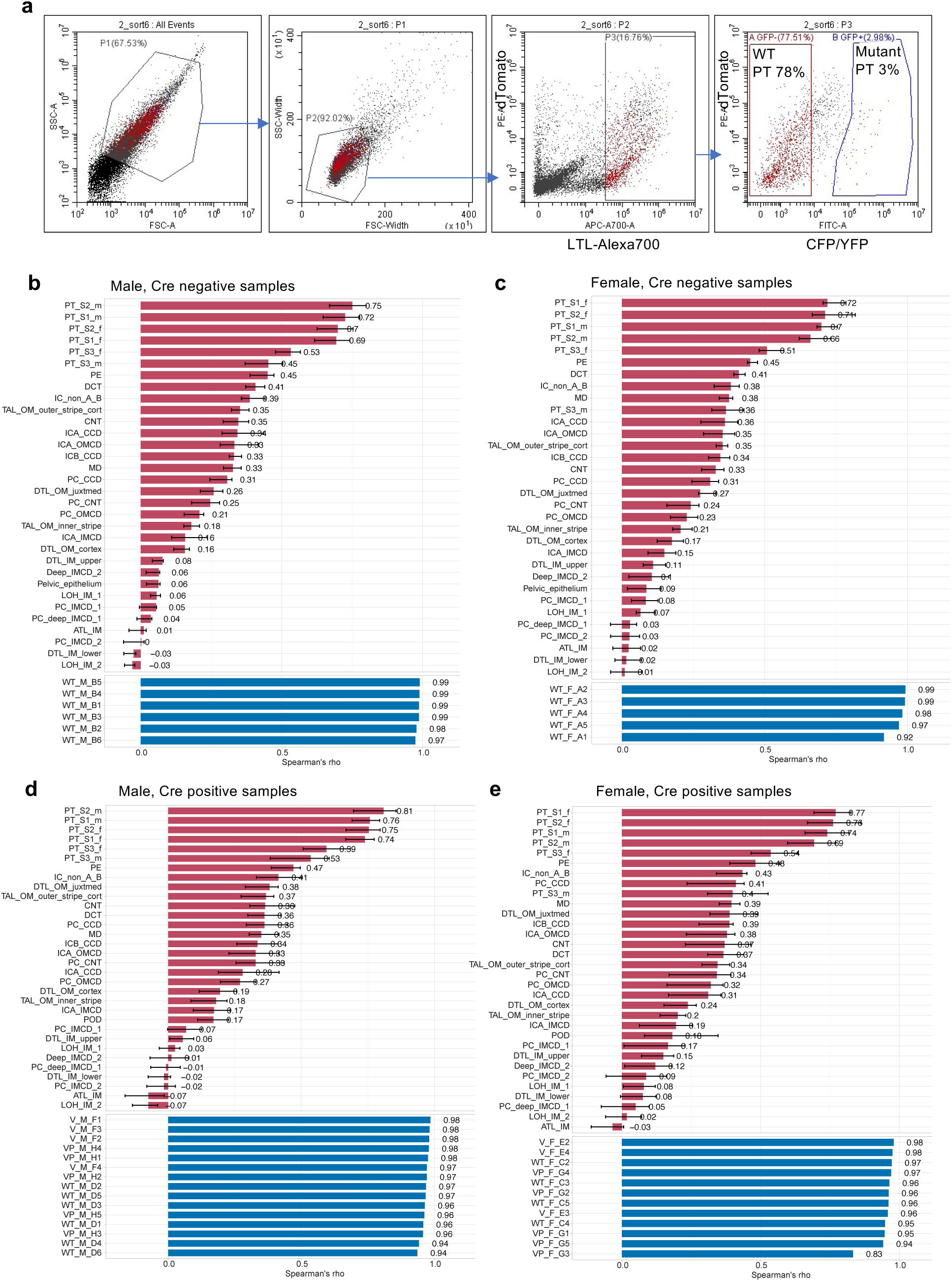
Isolation of renal proximal tubule cells for RNA-seq. **a** Representative flow cytometric sequential gating strategy to isolate single cells based on forward scatter and side scatter area and width, followed by gating for proximal tubule cells (positivity for LTL coupled to Alexa700) and then for negativity (no Cre activity) or positivity (Cre activity) for CFP/YFP. **b-e** CellMatchR algorithm analyses of the correlation of the overall transcriptomes of experimental samples to gene sets that are specific for individual renal epithelial cell types. Red bars show the mean ± std. dev. Spearman’s rho correlation of the test samples to each cell type signature. Blue bars show the inter-sample Spearman’s rho correlation of the test samples to one another. **b,c** Wild type proximal tubules without Cre activity isolated from male and female mice. **d,e** Proximal tubules with Cre activity isolated from male and female WT, *Vhl*, *Vhl/Pbrm1* mice. All epithelial cell type abbreviations are described in ^39^. For example, PT denotes proximal tubule, _S1, _S2 and _S3 denote S1, S2 and S3 segments, and _m or _f denote male or female. i.e. PT_S1_m refers to proximal tubule S1 segment male signature.

**Supplementary Fig. 4.**
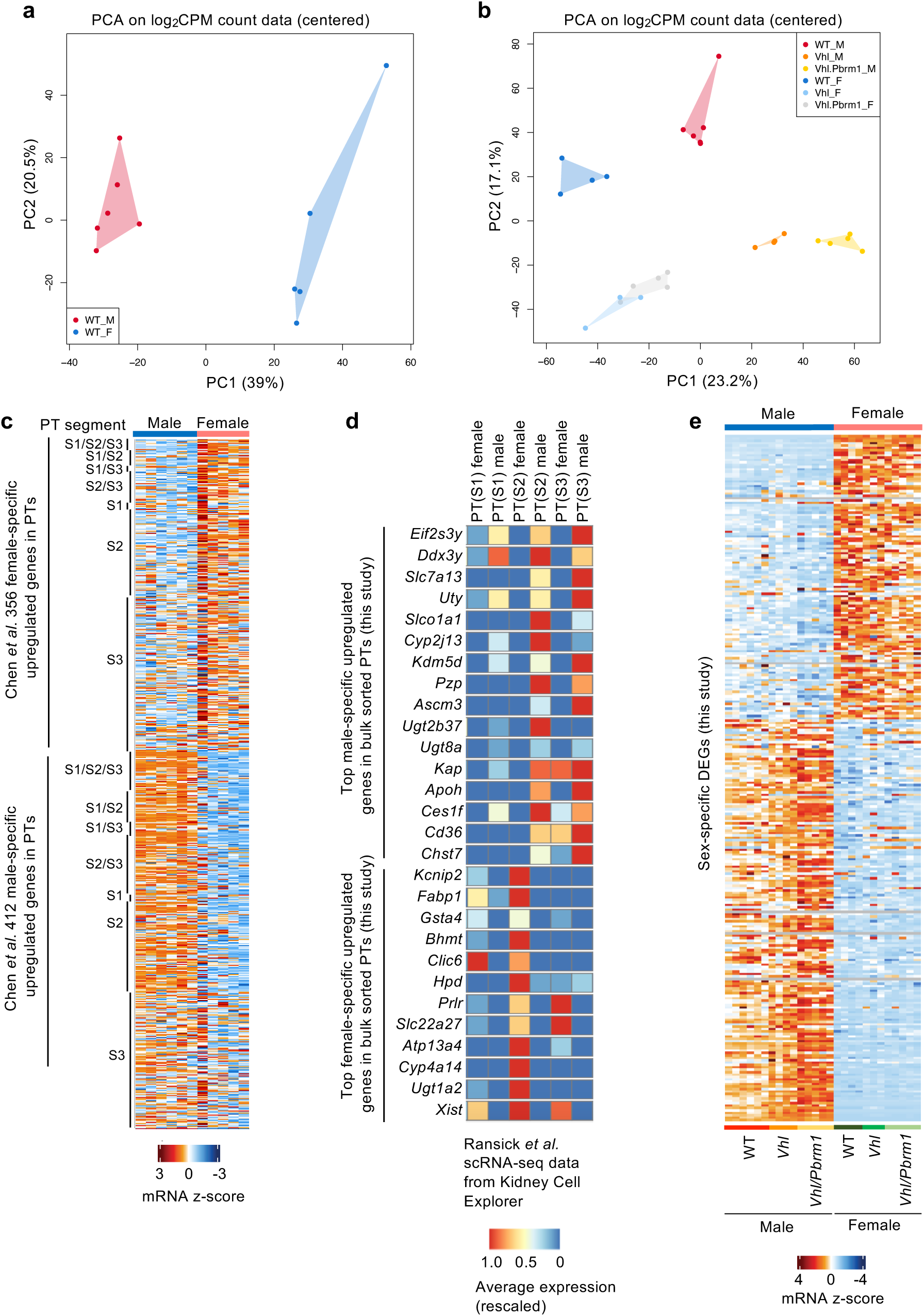
Analyses of sex-specific patterns of gene expression in proximal tubules. **a** Principal components analysis (log2CPM raw counts from RNA-seq) of proximal tubule cells without Cre activity isolated from male and female WT mice. **b** Principal components analysis (log2CPM raw counts from RNA-seq) of proximal tubule cells with Cre activity isolated from male and female WT, *Vhl*, *Vhl/Pbrm1* mutant mice. **c** Gene expression heatmap in our bulk RNA-seq of male and female proximal tubule cells probing for genes that were identified as being sex-specifically upregulated in S1, S2 or S3 proximal tubules (or in multiple combinations of these segments) in an snRNA-seq study ^6^. **d** Gene expression heatmap of the top sex-specific genes from our bulk RNA-seq of male and female proximal tubule cells used to probe a previously published scRNA-seq dataset ^8^ to compare relative expression levels in male and female S1, S2 and S3 proximal tubule cells. Data was obtained using Kidney Cell Explorer ^8^. **e** Gene expression heatmap of the sex-specific genes from Fig. 2a in proximal tubule cells obtained from the indicated genotypes and sexes. For all heatmaps the gene lists and their expression values are provided in the Source Data Table.

**Supplementary Fig. 5.**
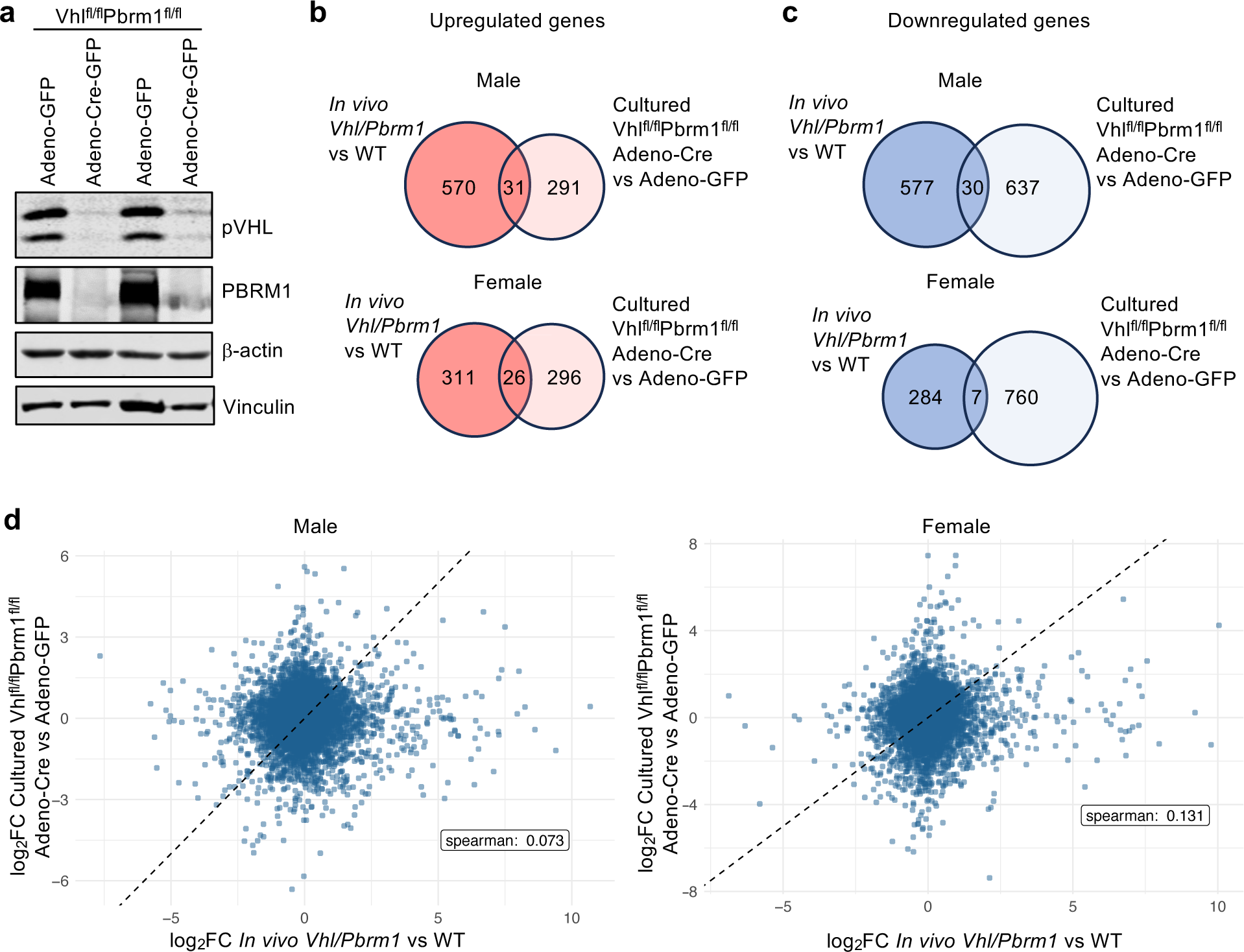
Transcriptomic effects of *Vhl/Pbrm1* deletion in cultured primary renal epithelial cells do not reflect the transcriptomic effects of *in vivo Vhl/Pbrm1* deletion. **a** Western blot demonstrating Adeno-Cre-GFP-mediated reduction of pVHL and PBRM1 in *Vhl^fl/fl^Pbrm1^fl/fl^* cultured primary renal epithelial cells 4 days after infection. Adeno-GFP served as a control. **b,c** Venn diagrams showing overlap of upregulated (**b**) and downregulated (**c**) genes between *in vivo* deletion of *Vhl/Pbrm1* compared to wild type proximal tubule cells and deletion of *Vhl/Pbrm1* (Adeno-Cre) compared to control (Adeno-GFP) in cultured *Vhl^fl/fl^Pbrm1^fl/f^*^l^ primary renal epithelial cells 4 days after infection. Male and female sets of genes were analysed separately in each setting. Cell cultures were established from three mice of each sex to provide mRNA for RNA-seq. Differentially expressed genes in each setting were defined as log2fc >1 or <-1 and *padj* < 0.05. **d** Plots comparing the relative changes in gene expression in *Vhl/Pbrm1* knockout versus wild type proximal tubules (x-axis) and cultured primary renal epithelial cells (y-axis) in males and females of all commonly-detected genes in the *in vivo* and cell culture datasets.

## Notes

**Conflict of interest statement:** The authors declare no potential conflicts of interest.

### Competing Interest Statement

The authors have declared no competing interest.

